# A multiscale model of the cardiovascular system that regulates arterial pressure via closed loop baroreflex control of chronotropism, cell-level contractility, and vascular tone

**DOI:** 10.1101/2021.10.21.465366

**Authors:** Hossein Sharifi, Charles K. Mann, Jonathan F. Wenk, Kenneth S. Campbell

## Abstract

Multiscale models of the cardiovascular system can provide new insights into physiological and pathological processes. PyMyoVent is a computer model that bridges from molecular to organ-level function and which simulates a left ventricle pumping blood through the systemic circulation. Initial work with PyMyoVent focused on the End Systolic Pressure Volume Relationship and ranked potential therapeutic strategies by their impact on contractility. This manuscript extends the PyMyoVent framework by adding closed loop feedback control of arterial pressure. The control algorithm mimics important features of the physiological baroreflex and was developed as part of a long-term program that focuses on growth and biological remodeling. Inspired by the underlying biology, the reflex algorithm uses an afferent signal derived from arterial pressure to drive a kinetic model that mimics the net result of neural processing in the medulla and cell-level responses to autonomic drive. The kinetic model outputs control signals that are constrained between limits that represent maximum parasympathetic and maximum sympathetic drive and which modulate heart rate, intracellular Ca^2+^ dynamics, the molecular-level function of both the thick and the thin myofilaments, and vascular tone. Simulations show that the algorithm can regulate mean arterial pressure at user-defined set-points as well as maintaining arterial pressure when challenged by changes in blood volume and/or valve resistance. The reflex also regulates arterial pressure when cell-level contractility is modulated to mimic the idealized impact of myotropes. These capabilities will be important for future work that uses computer modeling to investigate clinical conditions and treatments.

## Introduction

Multiscale models of the cardiovascular system can provide new insights into physiological and pathophysiological processes (Niederer et al. 2019a). Numerous groups are also trying to deploy them to improve the diagnosis and treatment of disease (Campbell et al. 2019; Niederer et al. 2019b). Models that incorporate molecular-level mechanisms may be particularly valuable for clinical applications because they can predict the effects induced by pharmaceutical therapies that alter the biophysical properties of molecules and/or the rate at which they undergo reactions.

Since cardiac output scales with stroke volume, multiscale models of the cardiovascular system often assume that changes in cell-level contractility produce corresponding changes in arterial pressure. *In vivo*, the situation is more complicated. In people, baroreceptors in the aortic arch and carotid sinuses fire action potentials at rates that vary with pressure. These signals are transmitted to the medulla which responds by modulating sympathetic and parasympathetic drive. This creates a feedback mechanism, termed the baroreflex, that adjusts chronotropism (heart rate), contractility, and vascular function to regulate arterial pressure (Beard et al. 2013; Jezek et al. 2022; Ursino and Magosso 2003).

The baroreflex operates continually, even at normal blood pressure, and modulates cell-level properties within seconds. Accordingly, the reflex provides short-term control, and is critical for stabilizing blood pressure after quick postural changes (Kaufmann et al. 2020). Other homeostatic mechanisms that regulate blood pressure, including the renin-angiotensin system (Harrison-Bernard 2009) and secretion of natriuretic peptides (Wong et al. 2017), primarily adjust renal excretion of salt and water, and consequently regulate blood pressure over much longer time-scales.

Since the baroreflex impacts multiple cell-level mechanisms, it can often compensate for an intervention that impacts a specific component of the cardiovascular system. As an example, if cell-level contractility is depressed with a negative myotrope such as mavacamten (Anderson et al. 2018; Green et al. 2016), the reflex is likely to up-regulate chronotropism, cell-level Ca^2+^-handling, and vascular tone to maintain arterial pressure. These reflex-mediated effects will change the contractile properties of the cells and the loads that they experience. In turn, this could change how the heart grows and adapts (Arts et al. 2005). As shown by Kerckhoffs et al. (2012) and Witzenburg & Holmes (Witzenburg and Holmes 2019), these reflex-mediated effects are important for simulating cardiac growth and ventricular remodeling.

This manuscript adds baroreflex control to the multiscale model previously described by Campbell et al. (2020). The goal was to mimic the key physiological effects that provide short-term regulation of arterial blood pressure. The algorithm is part of a larger framework that will be used in future work to simulate cardiac growth and biological remodeling. Accordingly, this paper focuses on the model’s response to perturbations and places less emphasis on clinical conditions that are best left for future work. In particular, the framework is explicitly not intended as a complete and/or fully detailed representation of the entire baroreflex. Other approaches including, but not restricted to, those developed by Beard et al. (Beard et al. 2013; Jezek et al. 2022; Kosinski et al. 2018), Olufsen et al. (Mahdi et al. 2013; Olufsen et al. 2006; Randall et al. 2019), and Ursino et al. (Silvani et al. 2011; Ursino 1998; Ursino and Magosso 2003) are more sophisticated and reproduce features, such as post-excitatory depression, that the current framework will not.

The current approach is based on a kinetic model that approximates the net result of neural processing in the medulla and signaling pathways in effector cells. The model is driven by an afferent signal derived from arterial pressure and outputs a control signal that is constrained between limits that represent maximum sympathetic and maximum parasympathetic drive. In turn, the control signal modulates heart rate, intracellular Ca^2+^ dynamics, the molecular-level function of both the thick and the thin filaments, and vascular tone. The results demonstrate that the algorithm can regulate mean arterial pressure at user-defined set-points, as well as maintaining arterial pressure when challenged by rapid changes in blood volume, valve resistance, and sarcomere-level contractility.

## Methods

### Overview

The model described here is called PyMyoVent. As originally described by Campbell et al. (2020), the framework uses a pacing stimulus to initiate a Ca^2+^ transient, that drives a myofilament-level model of contraction called MyoSim (Campbell 2014; Campbell et al. 2018). The contraction model predicts ventricular wall stress which is transformed into chamber pressure via Laplace’s law. Finally, the chamber pumps blood through a closed circulation mimicked using a series of resistances and compliant compartments. Together, these modules scale from molecular to organ-level function.

A prior publication (2020) investigated how module-level parameters (including ion channel permeabilities and myosin rate constants) affected system-level properties such as cardiac output. Heart rate was kept constant throughout these simulations. The current work extends the framework by implementing baroreflex control of arterial pressure (Figure 1). Specifically, heart rate and selected module level parameters were adjusted via negative feedback algorithms to drive arterial pressure towards a user-defined set-point. More details about the model for the Ca^2+^ transient and the baroreflex are provided immediately below. The other modules have already been described in detail (Campbell et al. 2020).

**Fig 1:**
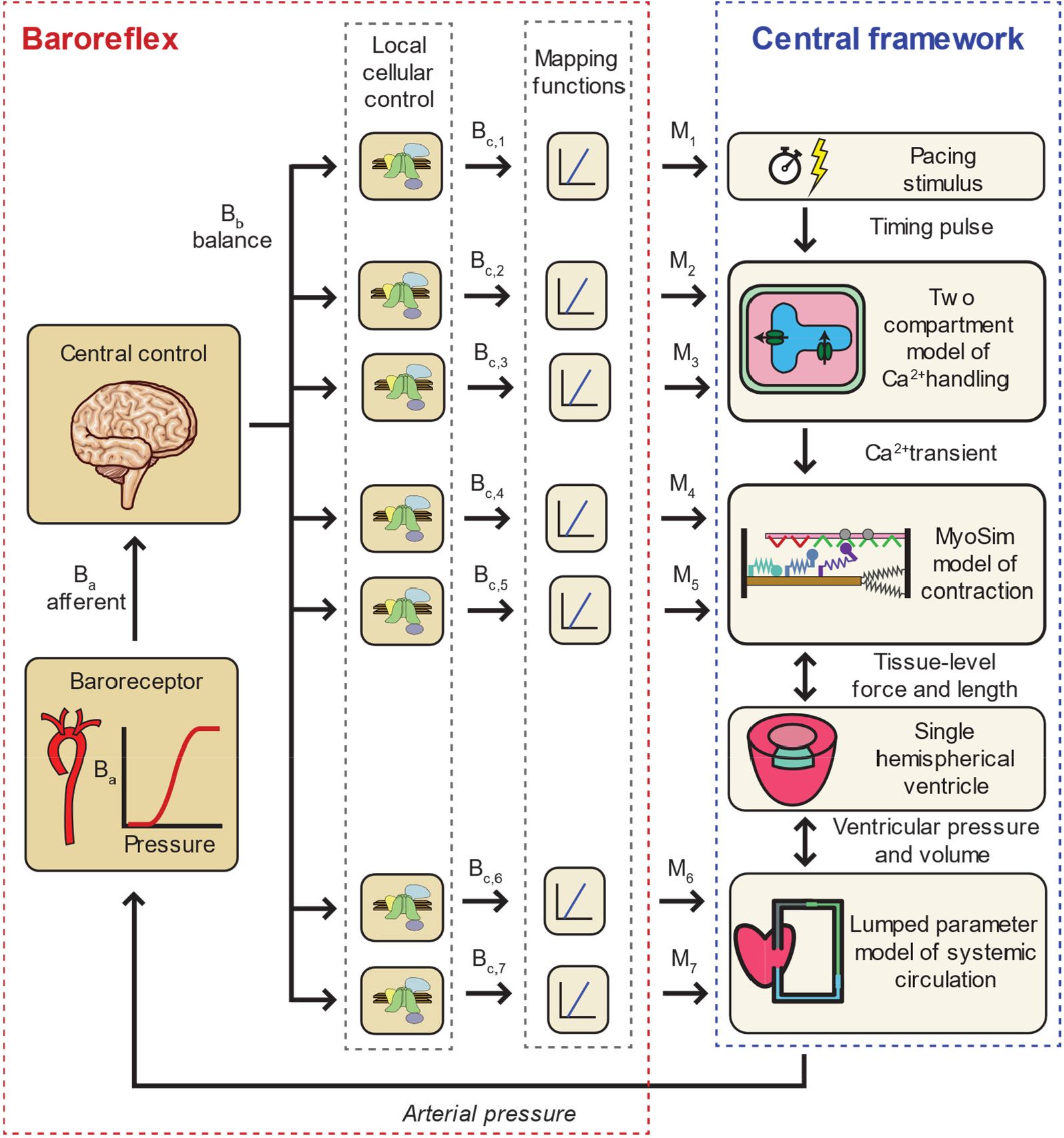
Overview of the PyMyoVent framework. The baroreflex algorithm monitors the arterial pressure predicted by the central framework and modulates heart rate, intracellular Ca^2+^ transients, myofilament contractility, and vascular tone in an attempt to drive the arterial pressure towards a user-defined setpoint. The B_a_, B_b_, B_c,i_ signals refer to the afferent, balance, and control signals while the M_i_ signals are the outputs of mapping functions. Further details are provided in the main text. Adapted from Campbell et al. (2020).

### Ca^2+^ transient

The team’s prior work used ten Tusscher et al.’s sophisticated model of myocyte electrophysiology (ten Tusscher et al. 2004) to simulate Ca^2+^ transients. Since most of the outputs from ten Tusscher et al.’s model were not required for the calculations, it was replaced here with a two compartment model that simplified the overall framework and increased the speed of the calculations. Specifically, the rates of change of the Ca^2+^ concentrations in the sarcoplasmic reticulum (Ca_SR_) and the myofilament space (Ca_myo_) were given by:

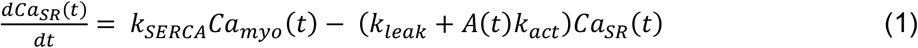

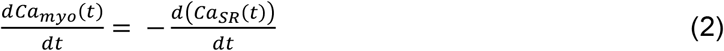

where the total Ca^2+^ concentration inside the cell Ca_total_ = Ca_SR_ + Ca_myo_ was kept constant, k_SERCA_ set the rate at which Ca^2+^ is pumped into the sarcoplasmic reticulum, k_leak_ defines a continual leak of Ca^2+^ into the myofilament space, and k_act_ controls the release of Ca^2+^ when the ryanodine receptors are open. A(t) is a pulse wave that is zero except for brief periods of duration t_open_ when A(t) is equal to one. These openings are initiated by the pacing stimuli that occur at time-intervals of t_RR_ and thus determine heart-rate.

Fig S1 in Supplementary Material shows that, with appropriately chosen parameters, the simple two compartment model simulates Ca^2+^ transients that are very similar to those calculated by Ten Tusscher et al.’s more sophisticated framework once the simulations have evolved to steady-state. Note that throughout this work, steady-state refers to a situation in which the envelopes of the calculated signals do not change with time.

### Baroreflex

Baroreflex control was implemented using an algorithm inspired by the underlying biology. Arterial pressure (P_arteries_) was transduced into a normalized afferent signal (B_a_) via the sigmoidal relationship

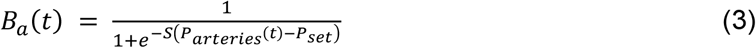

where P_set_ is the setpoint for arterial pressure, and S defines the slope of the function around its midpoint. B_a_ thus imitates the output of the baroreceptors and varies during the cardiac cycle due to the pulsatile nature of the arterial pressure signal.

In people, the medulla uses information encoded in the afferent signal to modulate the magnitudes of sympathetic and parasympathetic drive. These efferent signals regulate cellular-level processes in multiple organs so that sustained excess sympathetic drive increases arterial pressure and excess parasympathetic drive decreases it. The current model simplified these complex mechanisms using a single balance signal B_b_, seven distinct control signals (B_c,1_, B_c,2_ … B_c,7_), and seven mapping functions (M_1_, M_2_ … M_7_). As described in more detail below, the control signals and mapping functions modulated chronotropism, Ca^2+^ transients, myofilament function, and vascular tone.

The balance signal B_b_ is a normalized representation of the difference between sympathetic and parasympathetic efferent activity. Its rate of change was defined as

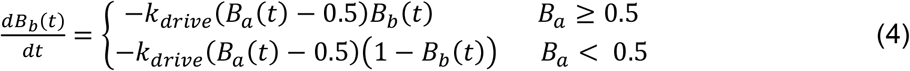

where k_drive_ is a rate constant. These equations cause B_b_ to tend towards one when sympathetic drive dominates the control loop and towards zero when parasympathetic drive predominates. k_drive_ sets the speed at which the control signal responds to changes in arterial pressure and/or P_set_.

The control signals B_c,i_ capture how each of the seven reflex-sensitive parameters in the cardiovascular model respond to the balance signal. Similar to equation 4, their rates of change were defined as

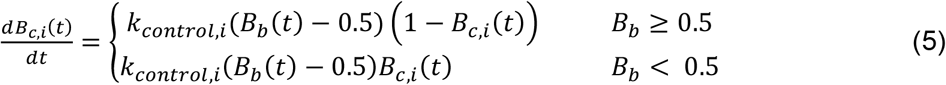

where i ranges from 1 to 7 and k_control,i_ is the rate constant for system i. These signals are also normalized and represent the status of cellular processes that are regulated by autonomic control. Each signal builds towards a saturating value of one when sympathetic drive exceeds parasympathetic drive (B_b_ > 0.5). If parasympathetic drive prevails, B_b_ is less than 0.5, and the control signals fall towards zero.

The final step in the algorithm used mapping functions M_i_ to link the normalized control signals B_c,I_ to actual parameter values. Each mapping function took the form

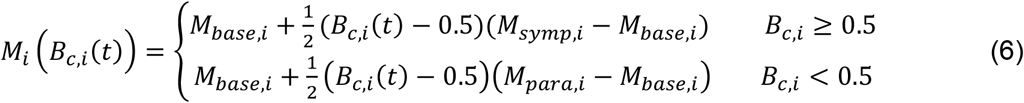

where M_base,i_ is the default value for parameter i, and M_symp,i_ and M_para,i_ are its limits during maximum sympathetic and maximum parasympathetic drive respectively.

Table 1 shows the mapping relationships. The M_base_, M_symp_ and M_para_ values for each parameter were defined in the model file (File S1 in Supplementary Material) that was used to initialize each simulation.

**Table 1:** Baroreflex implementation functions.

|  | Function | Maps to | Increased arterial pressure ... |
| --- | --- | --- | --- |
| Chrono-tropism | $M_1$ | $t_{\text{RR}}$ | Lengthens inter-beat interval and slows heart rate |
| Calcium-handling | $M_2, M_3$ | $k_{\text{SERCA}}$ and $k_{\text{act}}$ | Reduces the amplitude and prolongs the duration of $\text{Ca}^{2+}$ transients |
| Sarcomere contractility | $M_4, M_5$ | $k_1$ and $k_{\text{on}}$ | Reduces myosin cycling and sensitizes the thin filaments |
| Vascular tone | $M_6, M_7$ | $R_{\text{arteriole}}$ and $C_{\text{veins}}$ | Reduces systemic afterload and increases venous compliance |
| **Table 1: Baroreflex implementation functions** |  |  |  |

### Chronotropism

Baroreflex control of heart rate was implemented by mapping M_1_ to the inter-beat interval, t_RR_. The limits M_symp,1_ and M_para,1_ were set to 0.33 s and 1.5 s respectively so that heart-rate was constrained between 180 and 40 beats per minute.

### Cell-level contractility

Cell-level contractility was modulated using four parameters.

M_2_ and M_3_ mapped to k_act_ and k_SERCA_ (equation 1) respectively. The limits for these parameters were set so that increased arterial pressure reduced the amplitude and prolonged the duration of Ca^2+^ transients.

M_4_ and M_5_ modulated the k_1_ and k_on_ parameters in the MyoSim framework (Campbell et al. 2020; Campbell et al. 2018). The k_1_ parameter defines how quickly myosin heads transition from the super-relaxed (SRX) to the disordered relaxed (DRX) state (Schmid and Toepfer 2021) while k_on_ is the second-order rate constant for Ca^2+^-activation of binding sites on actin. Their limits (File S1) were set so that increased arterial pressure enhanced contractile force by biasing myosin heads towards the DRX state but desensitized the thin filament to Ca^2+^. These relationships mimic some of the effects produced by increased phosphorylation of myosin regulatory light chain and troponin I (Kampourakis et al. 2016; Solaro et al. 2013).

### Vascular tone

M_6_ and M_7_ controlled arteriolar resistance (R_arteriolar_) and venous compliance (C_veins_). File S1 shows that increased arterial pressure reduced R_arterioles_ and increased C_veins_. These effects reduce afterload and preload and complete the negative feedback loop.

### Implementation and computer code

Equations 4 and 5 were discretized and added to the system of ordinary differential equations that govern the PyMyoVent framework. The B_a_ signal was always calculated but the feedback was not applied (that is, B_b_ was not updated) until the reflex was activated in the code. This approach made it easier to discern how the reflex altered the system’s properties. Once the baroreflex had been activated, the inter-beat interval t_RR_ was updated once per cardiac cycle with all other feedback terms being updated on each time-step. This mimics the in vivo situation where the reflex provides continuous feedback control.

The code was written in Python using the Numpy (van der Walt et al. 2011), Scipy (Virtanen et al. 2020), and pandas (Reback et al. 2021) libraries. Full source code, examples, and scripts that reproduce the figures included in this manuscript are available at https://campbell-muscle-lab.github.io/PyMyoVent/. When run with a step-time of 0.5 ms, the calculations run ∼8 times slower than real-time on a single thread on a standard desktop PC. As an example, the simulation shown in Fig 2 took ∼30 minutes to complete.

**Fig 2:**
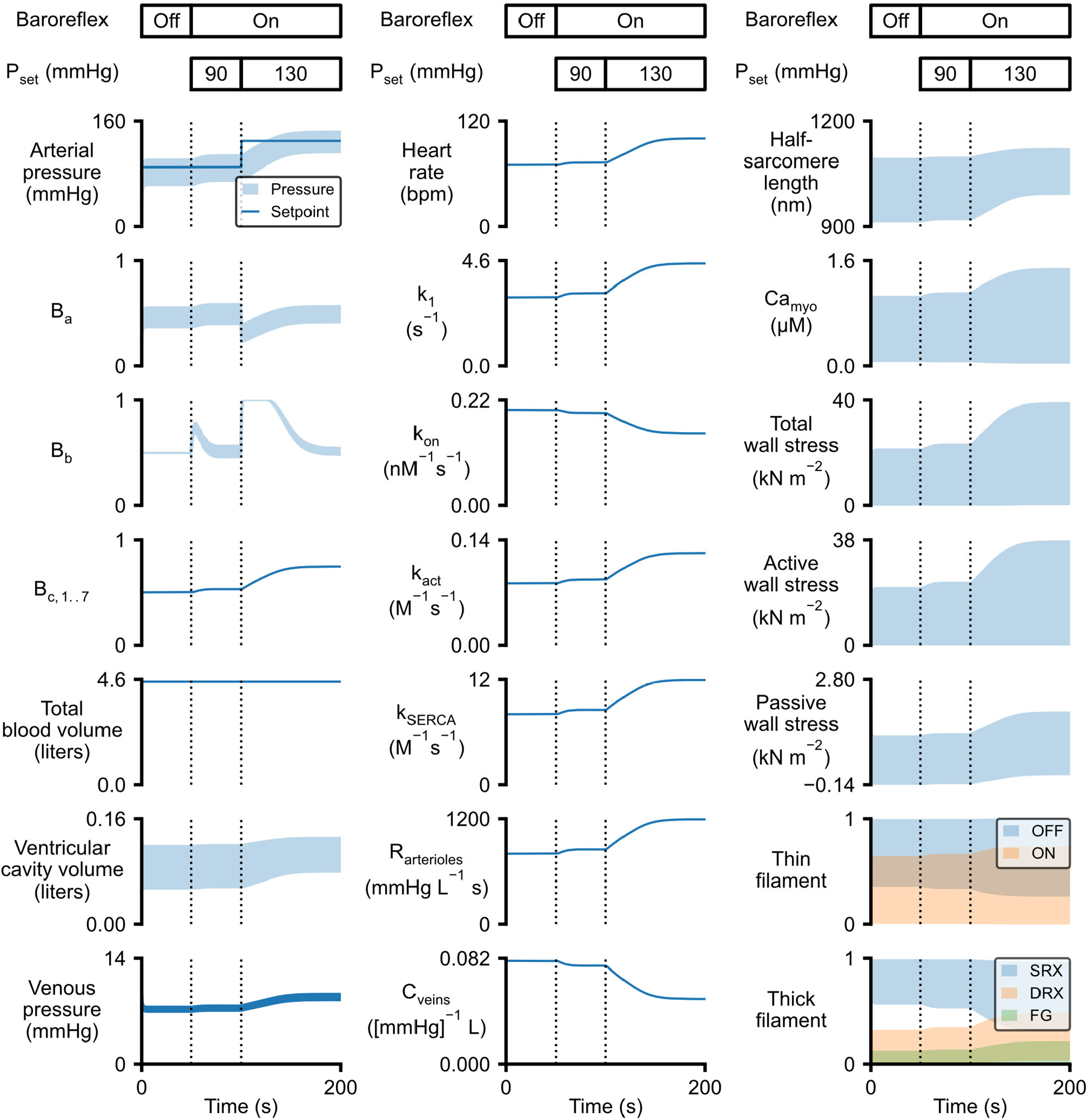
Simulation demonstrating baroreflex control of arterial pressure. The left hand column shows arterial pressure, the afferent (B_a_), balance (B_b_), and control (B_c,i_) signals for the baroreflex and 3 further system-level properties. The middle column shows the 7 parameters modulated by the baroreflex. The right-hand column shows properties relevant to myocardial function. The baroreflex was initiated after 50 s (first vertical line on each plot). The baroreflex setpoint was increased from 90 to 130 mmHg after 100 s (second vertical line on each plot). The OFF and ON labels describe the status of binding sites on the thin filament. The SRX, DRX, and FG labels refer to myosin heads in the super-relaxed, disordered relaxed, and force-generating states respectively (Campbell et al. 2020). The negative values for passive wall stress at end systole reflect the strain energy stored in the myocardium’s passive elastic structures (primarily titin and collagen) as active contraction shortens the myocytes below their resting length.

As described by Campbell et al. (2020), no attempt was made to constrain the parameters using experimental data. Instead, parameters were set to plausible values based on prior experience and typical values for a human. For example, the total blood volume for the systemic circulation was set to 4.5 liters. Similarly, for simplicity, the seven k_control,i_ parameters were set to the same value so that the B_c,i_ signals are identical in this work. The model could be enhanced in future work by setting different k_control,i_ values for each control. This would allow components of the reflex to respond at different speeds.

The default parameter values used to initialize all the simulations and the M_para,i_ and M_sump,i_ limits for each mapping function are shown in File S1. Some of these values differ from those in the original publication (Campbell et al. 2020) with the changes reflecting updates to the underlying code-base. Fig S2 shows the steady-state solution with these parameters for a single cardiac cycle. That figure is comparable to Fig 3 from Campbell et al. (2020) and is included here for completeness. The current work focuses on the baroreflex with most of the figures showing simulations that include at least 100 heart beats. In these figures, simulated values that change markedly during the cardiac cycle (for example, ventricular pressure) are shown as envelopes that outline the extreme values calculated for each heart-beat.

**Fig 3:**
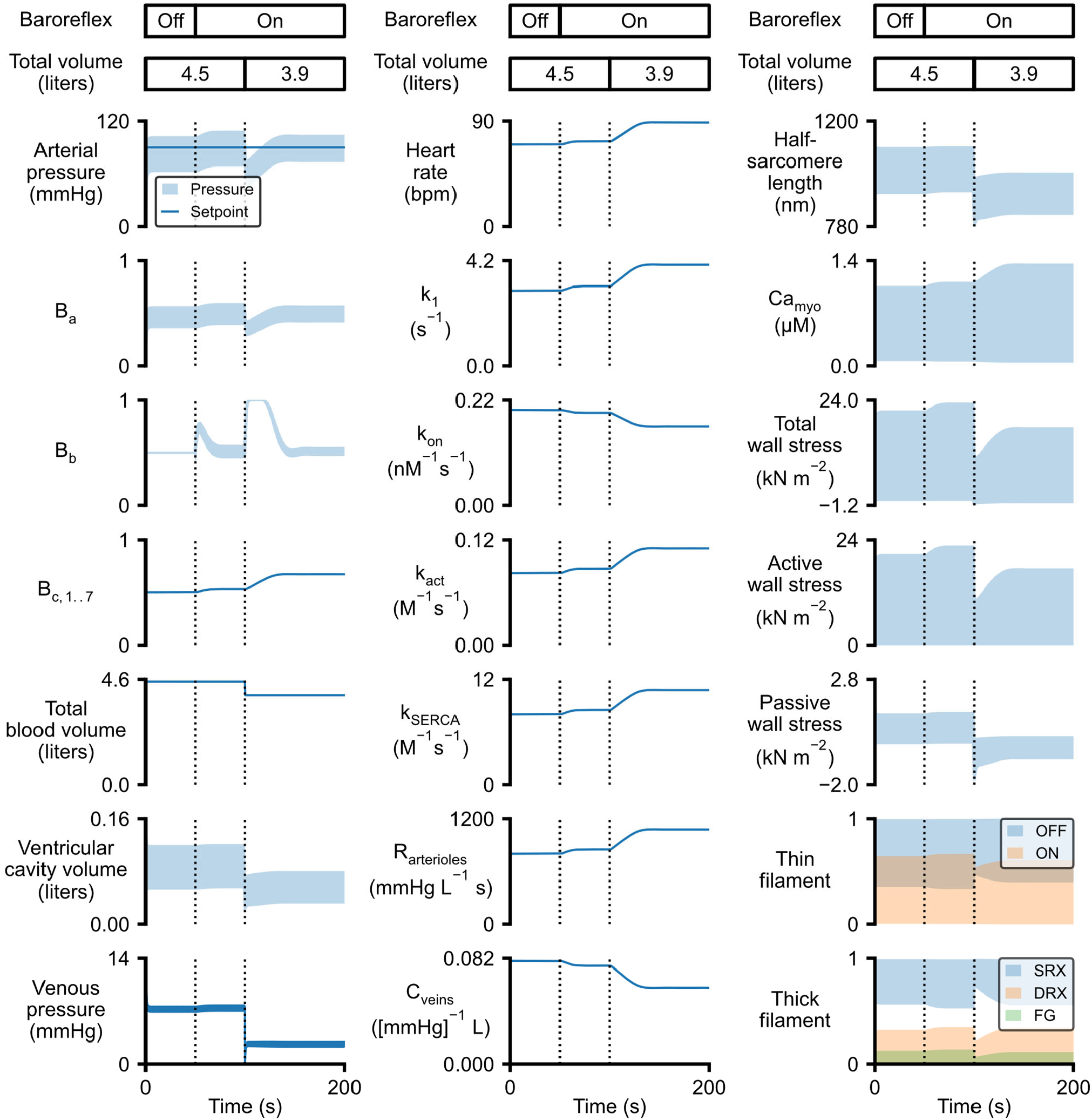
The baroreflex stabilizes arterial pressure after a reduction in blood volume. Figure panels are arranged as in Fig 2. The baroreflex was initiated after 50 s (first vertical line on each plot) with P_set_ equal to 90 mmHg. 600 ml of blood (∼13% of total volume) was removed from the venous circulation at 100 s (second vertical line). This perturbation induced rapid changes in heart rate, Ca^2+^-handling, myofilament contractility, and vascular tone which, in concert, maintained arterial pressure at the reflex setpoint. If the baroreflex had not been active, the blood loss would have reduced arterial pressure to 67/40 mmHg (see Fig S5).

## Results

### The baroreflex regulates arterial pressure at user-defined setpoints

Figure 2 demonstrates that the baroreflex algorithm implements closed loop control and regulates arterial pressure at user-defined setpoints. The simulation was initiated with the default parameters listed in File S1 and the reflex turned off in the code. The system reached steady-state (that is, the calculated values were stable) within 10 s at which time the mean arterial pressure was 83 mmHg and the mean B_a_ signal (calculated with P_set_ equal to 90 mmHg) was 0.47.

The baroreflex was activated at 50 s (the first dashed vertical line on each plot). Since the mean value of B_a_ was below its mid-point value of 0.5, the balance signal B_b_ started to increase (equation 4). This raised the B_c,i_ control signals (equation 5) and subsequently the M_i_ signals (equation 6) that regulated heart rate, myocyte Ca^2+^-handling, myofilament contractility, and vascular tone. These autonomic responses elevated arterial blood pressure and allowed the B_a_ afferent and the B_b_ balance signals to stabilize around their mid-points. Arterial pressure reached the new steady-state determined by the P_set_ value within ∼10 s of the reflex being activated.

The second dashed line in each plot at 100 s marks the time at which P_set_ was increased to 130 mmHg. This change induced a larger system-level response with the B_c,i_ control signals increasing by 0.21 as opposed to the 0.03 change that occurred when the reflex was activated. The B_c,I_ signals represent the status of cellular processes that are regulated by autonomic control with the signal values being determined by kinetic models (equation 5) that mimic the underlying biology. Since the speeds at which the B_c,i_ signals change are limited by the k_control,i_ rate constants, it required ∼30 seconds for the system to stabilize arterial pressure at the much higher set-point.

In summary, Figure 2 shows that the reflex algorithm can adjust system level properties to regulate arterial pressures at different levels. The responses are complete within tens of seconds and are sufficient to provide continual short-term feedback control.

Additional simulations were run with different values of P_set._ Fig S3 in Supplementary Material shows a simulation where arterial pressure was lowered to a set-point of 50 mmHg. These calculations demonstrate that the reflex could also reduce systemic pressure by reversing the physiological effects demonstrated in Fig 2.

Further tests (not shown) demonstrated that the reflex could regulate arterial pressure with P_set_ values ranging from ∼30 to ∼150 mmHg. Since this range is much broader than that linked to good cardiovascular health (roughly 65 to 100 mmHg) the algorithm is likely to be applicable to a wide range of conditions.

Fig S4 in Supplementary Material shows an example in which P_set_ was deliberately increased beyond the working range to 200 mmHg. This figure illustrates how the reflex algorithm responds to a perturbation that it cannot accommodate. The B_b_ balance signal increased to its maximum value of one but the controlled parameters saturated at the limits corresponding to maximum sympathetic drive without pressure reaching the setpoint. Note that a mean arterial pressure of 200 mmHg indicates a severe, and potentially fatal, hypertensive crisis and is far above the normal physiological range.

### The baroreflex stabilizes arterial pressure after a reduction in blood volume

Fig 3 shows the first of three simulations that demonstrate how the reflex maintains arterial pressure at a fixed set-point after different types of perturbation. In this first example, 600 ml of blood (∼13% of the total blood volume) was removed from the venous compartment while the reflex was active. Venous pressure dropped from 7 mmHg to 3 mmHg but the baroreflex modulated heart rate, Ca^2+^-transients, myofilament contractility, and vascular tone to maintain mean arterial pressure at the P_set_ value of 90 mmHg.

Without reflex control, the reduction in blood volume dropped mean arterial pressure to 54 mmHg (Fig S5 in Supplementary Material). Fig S6 in Supplementary Material superposes results from simulations with and without baroreflex control and emphasizes the different effects induced by the reduction in blood volume reduction and the reflex response.

Together, these figures demonstrate that the reflex algorithm can stabilize arterial pressure after blood volume is decreased.

### The baroreflex stabilizes arterial pressure when aortic resistance is increased

Fig 4 shows the system’s response to a sudden increase in aortic resistance. The perturbation increased the pressure gradient between the ventricle and the aorta and thus the afterload that the chamber ejected against. The myocytes shortened more slowly and stroke volume fell. Arterial pressure dropped as cardiac output diminished. The baroreflex responded to the sudden reduction in the B_a_ afferent by adjusting the B_b_ balance and B_c,i_ control signals to return arterial pressure back to the set-point.

**Fig 4:**
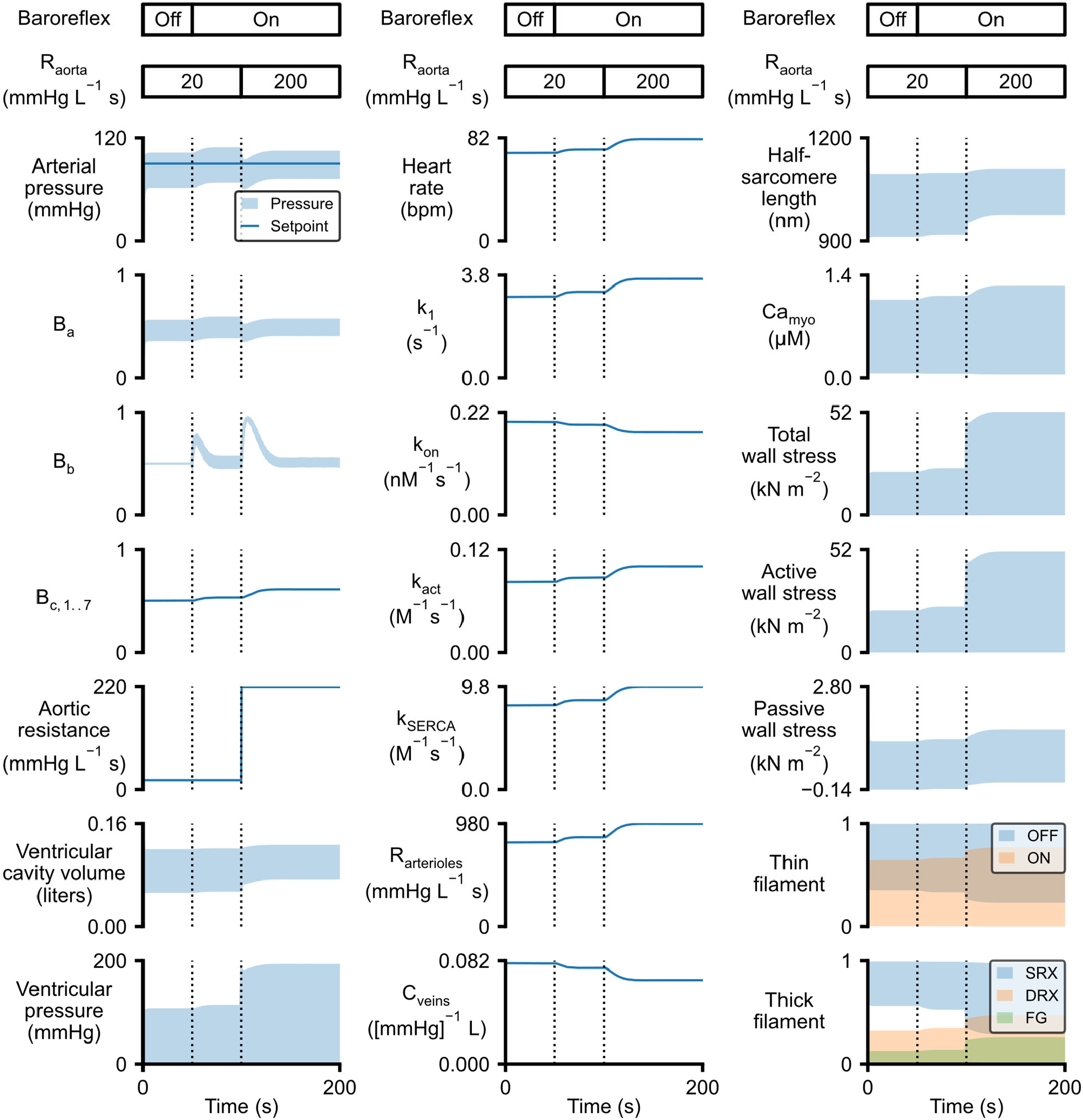
The baroreflex stabilizes arterial pressure when aortic resistance is increased. Figure panels are arranged as in Fig 2 except that aortic resistance is shown in place of total blood volume and ventricular pressure is shown in place of venous pressure. The baroreflex was initiated after 50 s (first vertical line on each plot) with P_set_ equal to 90 mmHg. The aortic resistance was increased from 20 to 220 mmHg L^-1^ s at 100 s. This perturbation induced changes in heart rate, Ca^2+^-handling, myofilament contractility, and vascular tone which together maintained arterial pressure at the reflex setpoint. Peak wall stress approximately doubled following the increase in aortic resistance. Fig S7 shows the system’s response to the same increase in aortic resistance without baroreflex control.

Results from an equivalent simulation performed without baroreflex control are shown in Fig S7 in Supplementary Material and compared to the reflex-controlled records in Fig S8. PV loops from the paired simulations are shown in Fig 5. These figures emphasize that increasing aortic resistance increases ventricular pressure irrespective of reflex control. (This is because the myocytes shorten more slowly and operate on a different portion of their force-velocity curve.) However, peak ventricular pressure is higher in the presence of reflex control because the baroreflex adjusts autonomic drive in order to maintain arterial pressure.

**Fig 5:**
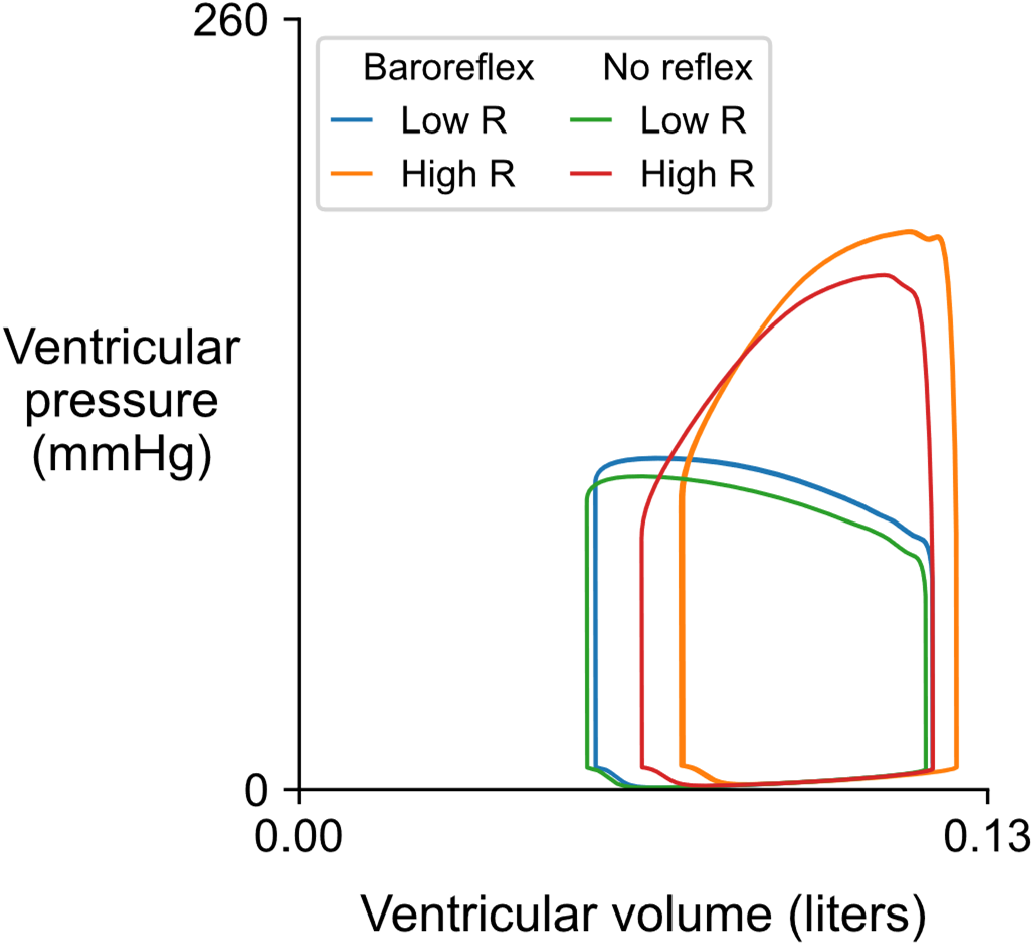
Pressure volume loops for a ventricle pumping against control and elevated aortic resistance with and without baroreflex control. The loops show simulated data from Figs 4 and S7 with ventricular pressure plotted against ventricular volume before and after the increase in aortic resistance (R). Note that ventricular pressure increases markedly even in the absence of baroreflex control as the myocardium ejects blood against the higher resistance. The reflex accentuates this response with the increased heart rate (not visible in this plot) acting to pump more blood into the aorta. Note that the decrease in vascular compliance induced by the reflex elevates filling pressure and thus end diastolic volume.

### The baroreflex stabilizes arterial pressure when myofilament contractility is reduced

Throughout this work, myofilament contraction was modeled using the MyoSim framework to simulate the dynamic behavior of cross-bridges cycling between super-relaxed (SRX), disordered-relaxed (DRX), and force-generating states (Campbell et al. 2018). The transition between the disordered-relaxed and super-relaxed states is governed by the k_2_ parameter. Increasing the value of this parameter biases heads towards the super-relaxed state and reduces contractility. This is likely to be one of the mechanisms through which the cardiac myotrope mavacamten reduces systolic force (Anderson et al. 2018).

Fig 6 shows that the reflex can regulate arterial pressure after myofilament contractility is perturbed by increasing the k_2_ parameter. Note that the responses include increases in heart rate, peak intracellular Ca^2+^ concentration, and vascular tone. As for the preceding perturbations, Fig S9 and S10 in Supplementary Material show how system-level function would have been affected if the reflex was not active.

**Fig 6:**
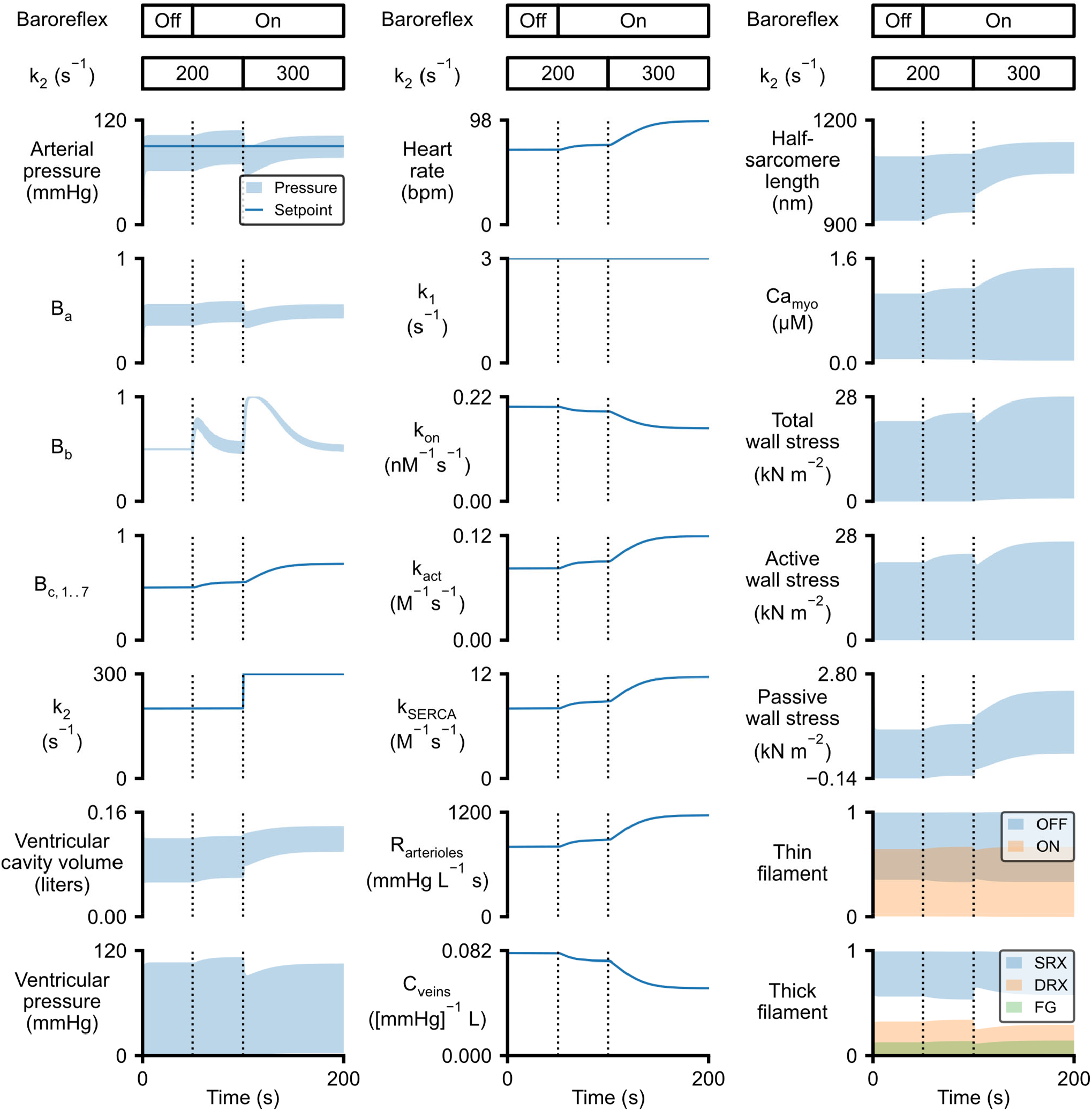
The baroreflex stabilizes arterial pressure when myofilament contractility is reduced. Figure panels are arranged as in Fig 4 except that the k_2_ parameter is shown in place of total blood volume. Increasing this parameter biases myosin heads towards the SRX state and reduces myofilament-level contractility. The baroreflex was initiated after 50 s (first vertical line on each plot) with P_set_ equal to 90 mmHg. k_2_ was increased from 200 to 300 s^-1^ at 100 s. This perturbation induced changes in heart rate, Ca^2+^-handling, myofilament contractility, and vascular tone which together maintained arterial pressure at the reflex setpoint. Fig S7 shows the system’s response to the same perturbation without baroreflex control.

### Baroreflex control of vascular tone contributes to homeostasis

Fig 8 shows a simulation similar to the example from Fig 2 but without the vascular components of the reflex; note that the values of R_arterioles_ and C_veins_ (bottom two panels in center column) remain at their defaults throughout the calculations.

**Fig 8:**
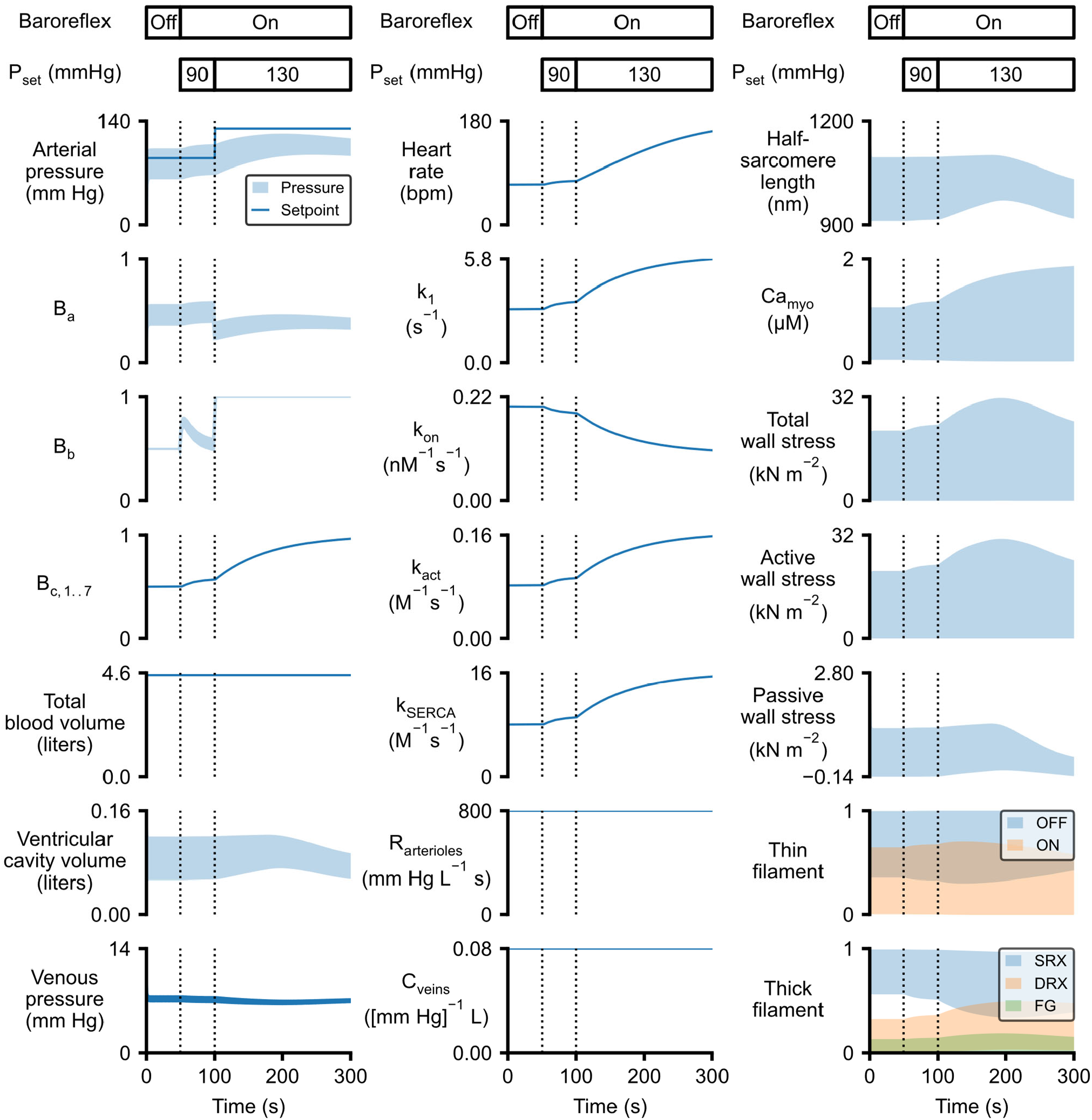
Simulation without baroreflex control of vascular tone. Figure panels are arranged as in Fig 2. The baroreflex was initiated after 50 s (first vertical line on each plot) with P_set_ being raised from 90 to 130 mmHg after 100s (second vertical line). This perturbation caused the reflex balance signal B_b_ to increase towards 1 and drove corresponding changes in heart rate, intracellular Ca^2+^-handling, and myofilament contractility. Arteriolar resistance and venous compliance were kept constant. Consequently, venous pressure remained roughly stable and ventricular filling (shown by cavity volume (left-hand column) and half-sarcomere relaxation (shown by the reduction in end-diastolic length in right-hand column) became impaired as heart rate increased.

When reflex control was activated (first dashed line in each panel in Fig 8), the adjustments to heart rate and contractility were sufficient to stabilize arterial pressure at the initial set-point of 90 mmHg. After 100 s (second dashed line in each panel), P_set_ was raised to 130 mmHg. This perturbation induced further increases in heart rate and contractility, and arterial pressure started to rise. However, unlike the situation in Fig 2 with the full reflex, venous compliance remained fixed so venous pressure was almost unchanged. As heart rate accelerated, the fixed venous pressure limited filling and end-diastolic volume began to fall. In turn, this compromised systolic function via Starling’s law (see active wall stress in third column of Fig 8) and arterial pressure declined. This simulation demonstrates that vascular tone is an important component of the baroreflex.

Fig S7 in Supplementary Material superposes data from Figs 2 and 8 in the main text to emphasize the effect of the vascular reflex.

## Discussion

This manuscript describes a mathematical model of the cardiovascular system that implements reflex control of arterial pressure by modulating heart rate, intracellular Ca^2+^-handling, myofilament biophysics, and vascular tone. The control algorithm was inspired by the underlying biology and incorporates several of the molecular-level mechanisms that are known to be regulated by the autonomic nervous system. Calculations showed that the framework could maintain arterial pressure at user-defined set-points within the physiological range, and at a fixed value when challenged by sudden changes in total blood volume, aortic resistance, or myofilament contractility.

The simulations also demonstrated the importance of dynamic control of vascular tone. The baroreflex’s ability to regulate arterial pressure was compromised when it could not adjust venous compliance in concert with heart-rate and cell-level contractility.

### Scope of this work

The goal of this work was to extend the PyMyoVent framework so that it can provide short-term regulation of arterial pressure after perturbations. This will be important for future work that will use PyMyoVent to simulate cardiac growth and biological remodeling because Kerchoffs (Kerckhoffs et al. 2012) and Witzenberg and Holmes (Witzenburg and Holmes 2019) have shown that the arterial pressure has an important impact on these processes. This issue was also discussed in a recent review (Sharifi et al. 2021).

In particular, it is important to emphasize that the current control algorithm is not intended to reproduce every feature of the real baroreflex. That would require a much more sophisticated approach, perhaps building on the specialized models developed by researchers including, but not restricted to, Beard, Ursino, and Olufsen (Jezek et al. 2022; Randall et al. 2019; Silvani et al. 2011).

Similarly, the perturbations shown in Figs 3 to 6 need to be interpreted with caution. While it is tempting to think of a reduction in blood volume as representing hemorrhage, an increase in aortic resistance as mimicking valve stenosis, and reduced contractility as equivalent to treatment with a myotrope, the real physiological situations are much more nuanced. One trivial difference is that the perturbations were imposed as step changes in the simulations rather than evolving with an appropriate time-scale. Valve conditions, for instance, often develop progressively over months and years. Another distinction is that the current framework only includes short-term regulatory mechanisms and does not adjust total blood volume. Adjustment of renal function and total blood volume is critical for long-term blood pressure control but is not yet represented in the model.

For these reasons, the current work focuses primarily on the algorithm and its mechanistic consequences. Discussions of clinical results and detailed comparisons with experimental data are best left for future studies that will build from this framework.

### Comparison with prior models incorporating baroreflex control

Many other mathematical models that incorporate baroreflex control have been published. Due to the complexity of the cardiovascular system, different groups have chosen to emphasize selected mechanisms and simplified other aspects of their model. A recent paper by Rondanina and Bovendeerd (2020), for example, focused on cardiac growth and implemented baroreflex control by adjusting only peripheral resistance and circulatory unstressed volume. Dupuis et al. (2018) used a similar approach but chose to adjust total blood volume rather than unstressed volume in their CircAdapt-based simulations (Palau-Caballero et al. 2016) of cardiac resynchronization therapy.

Other groups have added cardiac-based parameters to their baroreflex mechanisms. Jezek et al. (2021) investigated system-level responses to exercise and changes in posture, and adjusted heart rate, contractility, and vascular tone in their model. Ursino implemented similar mechanisms in their pioneering work (1998). Both of these papers idealized ventricle contraction using a time-varying elastance approach in which the relationship between ventricular pressure and volume varies with time during the cardiac cycle (Suga and Sagawa 1974).

The current model uses a different strategy that takes advantage of the molecular-level mechanisms underpinning the contractile module. Specifically, in addition to modulating heart rate and vascular tone, the baroreflex control algorithm adjusts four parameters that define the magnitude and time-course of intracellular Ca^2+^ transients and the function of the thick and thin myofilaments. To the authors’ knowledge, this is the first cardiovascular model in which the baroreflex algorithm directly controls molecular-level contractility rather than phenomenological relationships. This might make it easier to test how pharmaceuticals (as shown in Fig 6) and/or genetic modifications to calcium-handling proteins and/or sarcomeric proteins affect the baroreflex.

The kinetic model described by equations 3 to 5 is also interesting because it captures several aspects of the underlying physiology in a mathematical framework. These factors include the concept of excess autonomic drive which is represented in the model by the balance signal B_b_. If arterial pressure is below the reflex setpoint, the medulla increases sympathetic efferent activity and decreases parasympthatic activity. In the calculations, this is mimicked by B_b_ rising towards its upper limit of one. Similarly, arterial pressures greater than the reflex setpoint will increase parasympathetic activity and diminish sympathetic drive. The model mimics this behavior by driving B_b_ towards zero. The ordinary differential equation (equation 4) dictates how the B_b_ signal evolves over time with the rate constant k_drive_ controlling how quickly the signal responds to changes in arterial pressure and setpoint.

The B_c,i_ signals (equation 5) provide an additional level of control and are intended to mimic the response of the cell-level effectors to changing autonomic drive. B_c,5_, for instance, modulates the second order rate constant k_on_ for Ca^2+^ activation of binding sites on the thin filament (Campbell et al. 2018). In vivo, this is regulated, at least in part, by Protein Kinase A-based phosphorylation of troponin I (Solaro et al. 2013). Excess sympathetic drive enhances kinase activity and, if sustained, gradually increases the proportion of troponin I molecules that are phosphorylated. The value of k_on_ when all of the troponins are phosphorylated is the M_symp,5_ limit. Similarly, excess parasympathetic drive (B_b_ < 0.5) deactivates the kinase allowing the proportion of troponin I molecules to fall over time. The M_para,5_ limit defines the k_on_ value when none of the troponin molecules are phosphorylated. The other control signals mimic similar mechanisms. In all cases, the differential equations (equation 5) allow the status of the mechanisms to evolve over time with the speed of the responses being set by the k_control,i_ parameters.

### Importance of vascular tone

Prototype versions of the current model attempted to regulate arterial pressure solely via chronotropism and adjustments to cell-level contractility. As demonstrated in Figs 8, these attempts were only partly successful and failed to regulate arterial pressure over a realistic physiological range.

The problem with the prototypes was that venous return did not increase with cardiac output. When the reflex was activated, contractility and heart-rate increased to elevate cardiac output and raise arterial pressure. While this strategy was successful for the first few beats, it also redistributed blood towards the arteries. Since the veins remained compliant, the pressure difference pushing blood back towards the heart dropped slightly and venous return could not sustain the increased cardiac output. End-diastolic volume started to fall with consequent reductions in stroke volume. Heart rate continued to increase in an attempt to compensate but this further compromised filling and the cardiac ejection fell.

Adding reflex control of vascular tone solved this problem and allowed the algorithm to maintain arterial pressure over a wider operating range. The key addition was allowing venous compliance to fall during increased sympathetic drive. This raised venous pressure and enhanced venous return so that it could sustain long-term increases in cardiac output.

### Limitations of the current model

Some of the limitations of the PyMyoVent framework have already been discussed (Campbell et al. 2020). These include: the single ventricle architecture, the assumption that the ventricle is hemispherical, and the one-way coupling of the electrophysiological and contractile modules. Each of these factors leads to further consequences. For example, the single ventricle architecture eliminates the possibility of atrial kick and diastolic filling thus occurs in a single phase.

The baroreflex developed in this work is also greatly simplified. One of the most obvious issues is that real hearts receive dual input from both parasympathetic and sympathetic nerves while PyMyoVent uses a single B_b_ balance signal which reflects the efferents’ difference. One of the limitations of this approach is that it prevents adjusting single reflex effectors independently of the others.

Another weakness is that the current model only incorporates a few of the mechanisms that are subject to autonomic control. Using the myofilaments as an example, sympathetic drive leads to phosphorylation of, at a minimum, troponin I, troponin T, myosin regulatory light chain, myosin binding protein-C, and titin (Crocini and Gotthardt 2021). In turn, these posttranslational modifications converge to impact a few key properties, such as Ca^2+^-sensitivity. Capturing all of these molecular effects in a single model would be challenging and, even if it could be done - perhaps using a spatially explicit model such as FiberSim (Kosta et al. 2021) - the parameters quantifying each effect’s magnitude would be poorly constrained because of the confounding and overlapping pathways.

Accordingly, the current algorithm uses a different approach that approximates baroreflex control by modulating a small, and hopefully representative, number of mechanisms distributed across the different components. Sarcomeric function, for example, is regulated by just two pathways which modulate thick filament activation (Campbell et al. 2018) and thin filament Ca^2+^ sensitivity (Solaro et al. 2013) respectively. The algorithm prioritizes simplicity while retaining the ability to simulate the key effects underlying short-term regulation of arterial pressure.

### Summary

PyMyoVent is a multiscale model of the cardiovascular system that can regulate arterial pressure using a baroreflex algorithm that is inspired by the underlying biology and regulates molecular-level mechanisms. The reflex framework was developed as part of a long-term program to use computer models to help optimize therapies for patients who have heart failure. The software runs on a standard laptop and is fully open-source. Interested parties are encouraged to download the software and use it in their own work. The current authors hope to use the new system to improve quantitative understanding of cardiovascular function and potentially to help accelerate the development of better therapies for patients who have cardiovascular disease.

## Acknowledgements

Supported by NIH HL133359 and HL163977 to KSC and JFW, NIH 148785 and TR0001998 to KSC, and AHA TP135689 to KSC.

## Author contributions

HS drafted the manuscript, wrote prototype versions of the code, contributed to the PyMyoVent repository and website, and ran prototype simulations. KJM helped developed the model framework. JFW helped develop the model framework and edited the manuscript. KSC planned the overall project, developed the baroreflex algorithm, wrote the final version of the code, ran the final simulations, created the figures, and finalized the manuscript.

