## Supplementary material for "A multiscale model of the cardiovascular system that regulates arterial pressure via closed loop baroreflex control of chronotropism, cell-level contractility, and vascular tone"

Hossein Sharifi [1]

Charles K. Mann [1]

Jonathan F. Wenk [1,2]

Kenneth S. Campbell [3]

[1] Department of Mechanical Engineering, University of Kentucky, Lexington, KY

[2] Department of Surgery, University of Kentucky, Lexington, KY

[3] Department of Physiology and Division of Cardiovascular Medicine, University of Kentucky, Lexington, KY

File S1. Default parameters for the PyMyoVent simulations.

This file is written in JSON format and was used to initialize most of the simulations presented in this work. (Exceptions are explained in the text). The $M_{para,i}$ and $M_{symp,i}$ limits for each reflex-controlled parameter were defined using the para_factor and symp_factor variables for the appropriate control. As an example, using the first control in the baroreflex section, $t_{RR}$ = ($t_{active\_period}$ + $t_{quiescent\_period}$) was initialized at 0.857 s and constrained between a parasympathetic limit of para_factor x $t_{RR}$ = 1.754 x 0.857 s = 1.5 s (~40 beats per minute) and a sympathetic limit of symp_factor x $t_{RR}$ = 0.386 x 0.857 s = 0.331 s (~180 beats per minute).

Further details are available at https://campbell-muscle-lab.github.io/PyMyoVent/.


```
{
    "PyMyoVent”:{
      "version”: “2.1.0”
    },
    "circulation":{
        "blood_volume": 4.5,
        "compartments":
        [
            {
                "name": "aorta",
                "resistance": 20,
                "compliance": 4e-4,
                "slack_volume": 0.3,
                "inertance": 0
            },
            {
                "name": "arteries",
                "resistance": 20,
                "compliance": 8e-4,
                "slack_volume": 0.3,
                "inertance": 0
            },
            {
                "name": "arterioles",
                "resistance": 800,
                "compliance": 1e-3,
                "slack_volume": 0.1,
                "inertance": 0
            },
            {
                "name": "capillaries",
                "resistance": 150,
                "compliance": 1e-4,
                "slack_volume": 0.25,
                "inertance": 0
            },
            {
                "name": "venules",
                "resistance": 50,
                "compliance": 0.03,
                "slack_volume": 0.5,
                "inertance": 0
            },
            {
                "name": "veins",
                "resistance": 20,
                "compliance": 0.08,
                "slack_volume": 2.0,
                "inertance": 0
            },
            {
```

```
            "name": "ventricle",
            "resistance": 5,
            "slack_volume": 0.065,
            "wall_density": 1055,
            "wall_volume": 0.1,
            "inertance": 0
        }
    ]
},
"heart_rate": {
    "t_active_period": 0.003,
    "t_quiescent_period": 0.854,
    "t_first_activation": 0.1
},
"half_sarcomere":{
    "initial_hs_length": 900,
    "reference_hs_length": 1100,
    "membranes": {
        "Ca_content": 1e-3,
        "k_leak": 6e-4,
        "k_act": 8.2e-2,
        "k_serca": 8.0,
        "t_open": 0.01,
        "implementation":{
            "kinetic_scheme": "simple_2_compartment"
        }
    },
    "myofilaments":{
        "cb_number_density": 1.15e17,
        "prop_fibrosis": 0.0,
        "prop_myofilaments": 0.6,
        "k_1": 3,
        "k_force": 1e-3,
        "k_2": 200,
        "k_3": 32,
        "k_4_0": 30,
        "k_4_1": 1,
        "k_cb": 0.001,
        "x_ps": 5,
        "k_on": 2e8,
        "k_off": 200,
        "k_coop": 5,
        "int_passive_exp_sigma": 300,
        "int_passive_exp_L": 70,
        "int_passive_l_slack": 950,
        "ext_passive_exp_sigma": 300,
        "ext_passive_exp_L": 70,
        "ext_passive_l_slack": 950,
        "implementation":
        {
            "kinetic_scheme": "3_state_with_SRX",
            "int_passive_mode": "exponential",
            "ext_passive_mode": "exponential",
            "max_rate": 2000,
            "temperature": 310,
            "bin_min": -10,
            "bin_max": 10,
            "bin_width": 1,
            "filament_compliance_factor": 0.5,
            "thick_filament_length": 815,
            "thin_filament_length": 1120,
            "bare_zone_length": 80,
            "reference_hsl_0": 1100,
            "delta_G_ATP": 70000,
            "thick_wall_approximation": true
```

```
            }
        }
    },
    "baroreflex":
    {
        "baro_P_set": 90,
        "baro_S": 0.02,
        "baro_k_drive": 10,
        "baro_k_recov": 0.1,
        "controls":
        {
            "control":
            [
                {
                    "level": "heart_rate",
                    "variable": "t_quiescent_period",
                    "k_control": 0.025,
                    "k_recov": 0.1,
                    "para_factor": 1.753497,
                    "symp_factor": 0.386
                },
                {
                    "level": "membranes",
                    "variable": "k_act",
                    "k_control": 0.025,
                    "k_recov": 0.1,
                    "para_factor": 0.5,
                    "symp_factor": 2.0
                },
                {
                    "level": "membranes",
                    "variable": "k_serca",
                    "k_control": 0.025,
                    "k_recov": 0.1,
                    "para_factor": 0.5,
                    "symp_factor": 2
                },
                {
                    "level": "myofilaments",
                    "variable": "k_1",
                    "k_control": 0.025,
                    "k_recov": 0.1,
                    "para_factor": 0.5,
                    "symp_factor": 2
                },
                {
                    "level": "myofilaments",
                    "variable": "k_on",
                    "k_control": 0.025,
                    "k_recov": 0.1,
                    "para_factor": 2,
                    "symp_factor": 0.5
                },
                {
                    "level": "circulation",
                    "variable": "arterioles_resistance",
                    "k_control": 0.025,
                    "k_recov": 0.1,
                    "para_factor": 0.5,
                    "symp_factor": 2
                },
                {
                    "level": "circulation",
                    "variable": "veins_compliance",
                    "k_control": 0.025,
```

```
                    "k_recov": 0.1,
                    "para_factor": 4,
                    "symp_factor": 0.25
                }
            ]
        }
    }
}
```

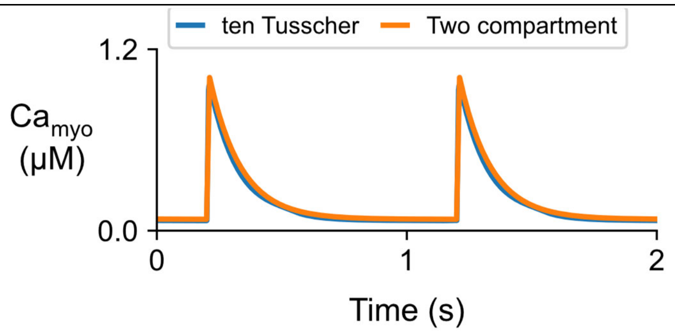


Fig S1: Comparison of $Ca^{2+}$ transients predicted by the ten Tusscher et al. and two-compartment models.

The blue trace (partially hidden) shows the steady-state intracellular $Ca^{2+}$ signal predicted by Ten Tusscher et al.'s model (2004) with default parameters. The orange trace shows the steady-state prediction for the two-compartment model (Equations 1 and 2 in main text) used in this work.

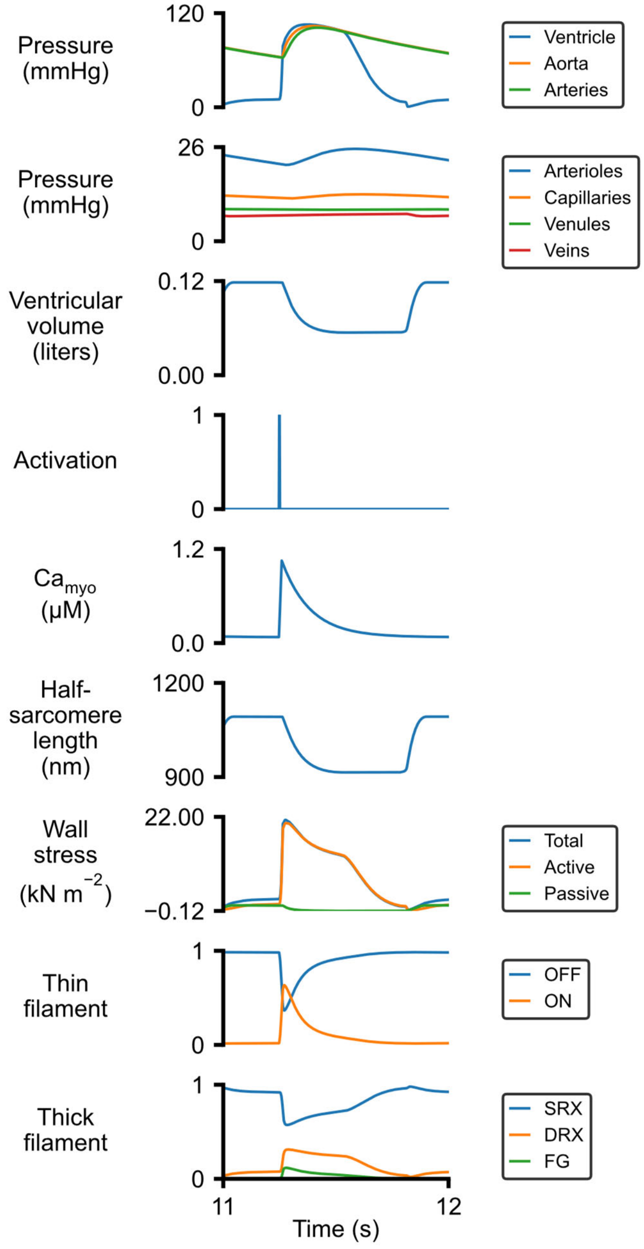


Fig S2: Steady-state cardiac cycle with default parameters.

The top 3 panels show system-level properties (pressures and ventricular volume) while the lower panels show signals related to calcium transients and myofilament function. The OFF and ON labels describe the status of binding sites on thin filament. The SRX, DRX, and FG labels refer to myosin heads in the super-relaxed, disordered-relaxed, and force-generating states respectively (Campbell et al. 2018).

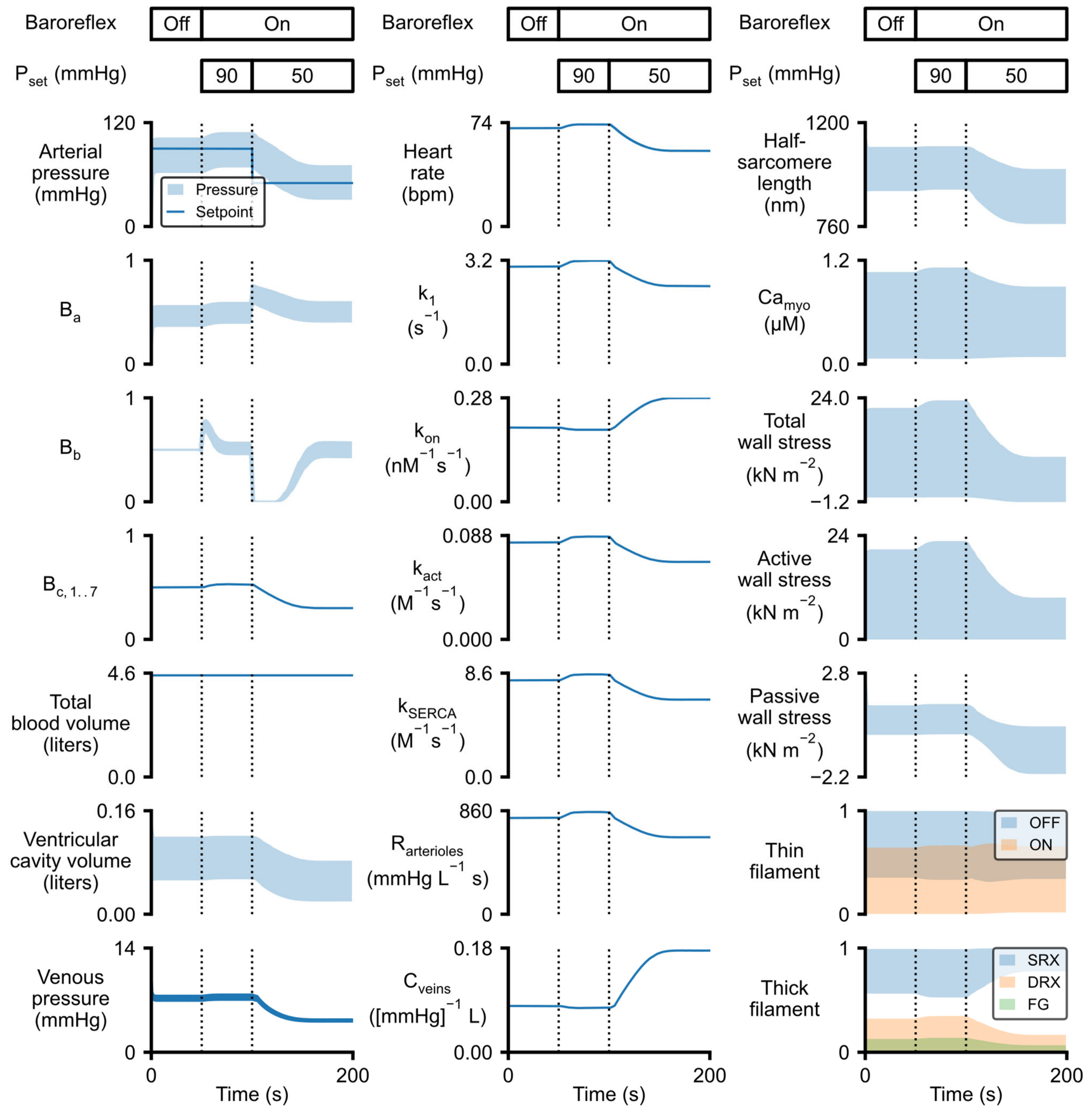


Fig S3: Baroreflex response to a reduction in the reflex setpoint.

This figure is identical to Fig 2 in the main text except that $P_{set}$ was reduced to 50 mmHg at 100 s (second vertical line on each plot).

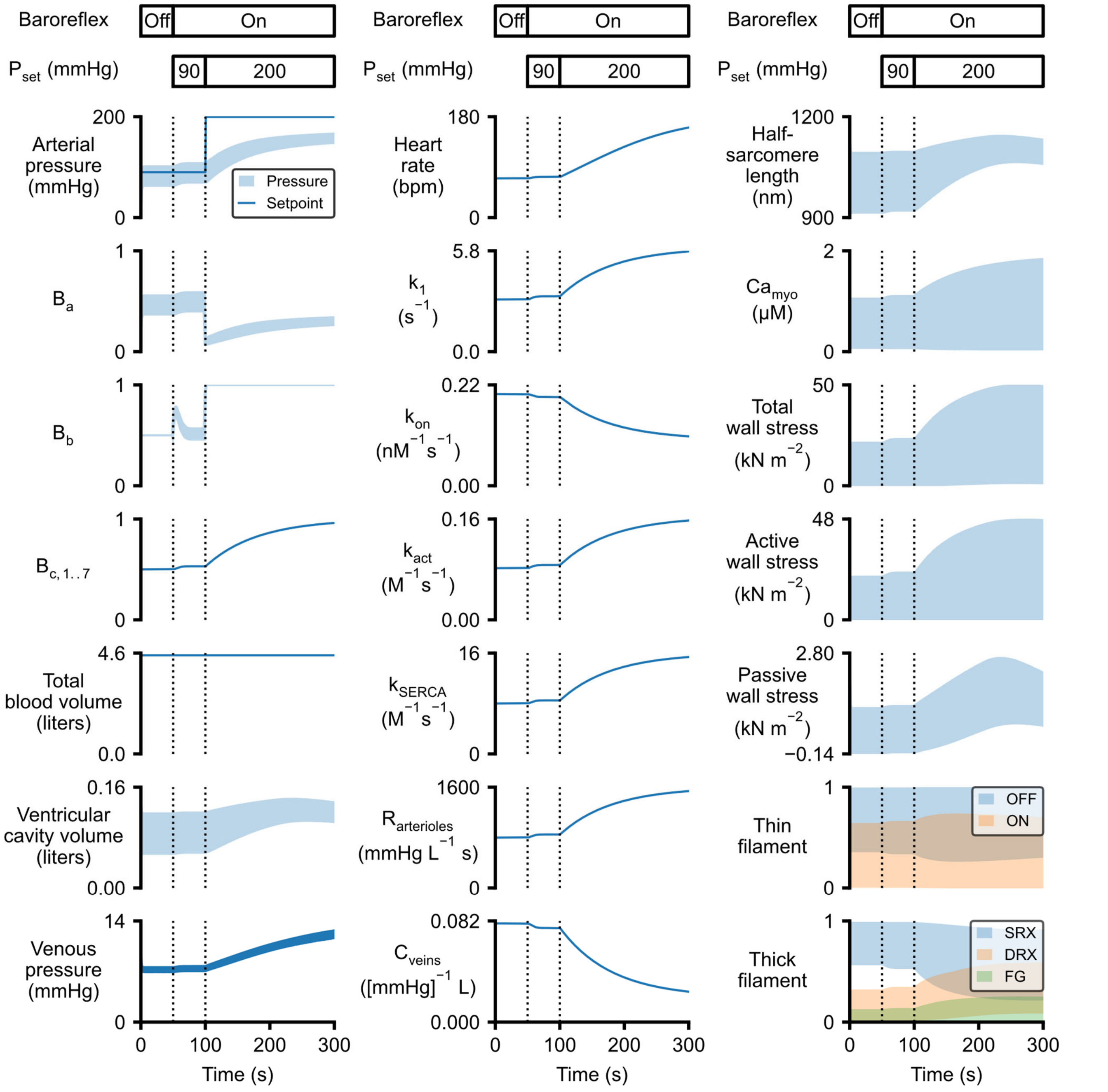


Fig S4: A simulation with $P_{set}$ raised above the operating range.

Figure panels are arranged as in Fig 2 in the main text. The baroreflex was initiated after 50 s (first vertical line on each plot) with $P_{set}$ being raised from 90 to 200 mmHg after 100s (second vertical line). This perturbation caused the balance signal $B_b$ to increase towards 1. This drove the control signals $B_{c,i}$ towards 1 so that the reflex-controlled mechanisms saturated at levels corresponding to maximum sympathetic drive. Peak arterial pressure rose to a maximum value of 172 mmHg but the reflex -controlled responses were insufficient to raise arterial pressure to the $P_{set}$ value of 200 mmHg.

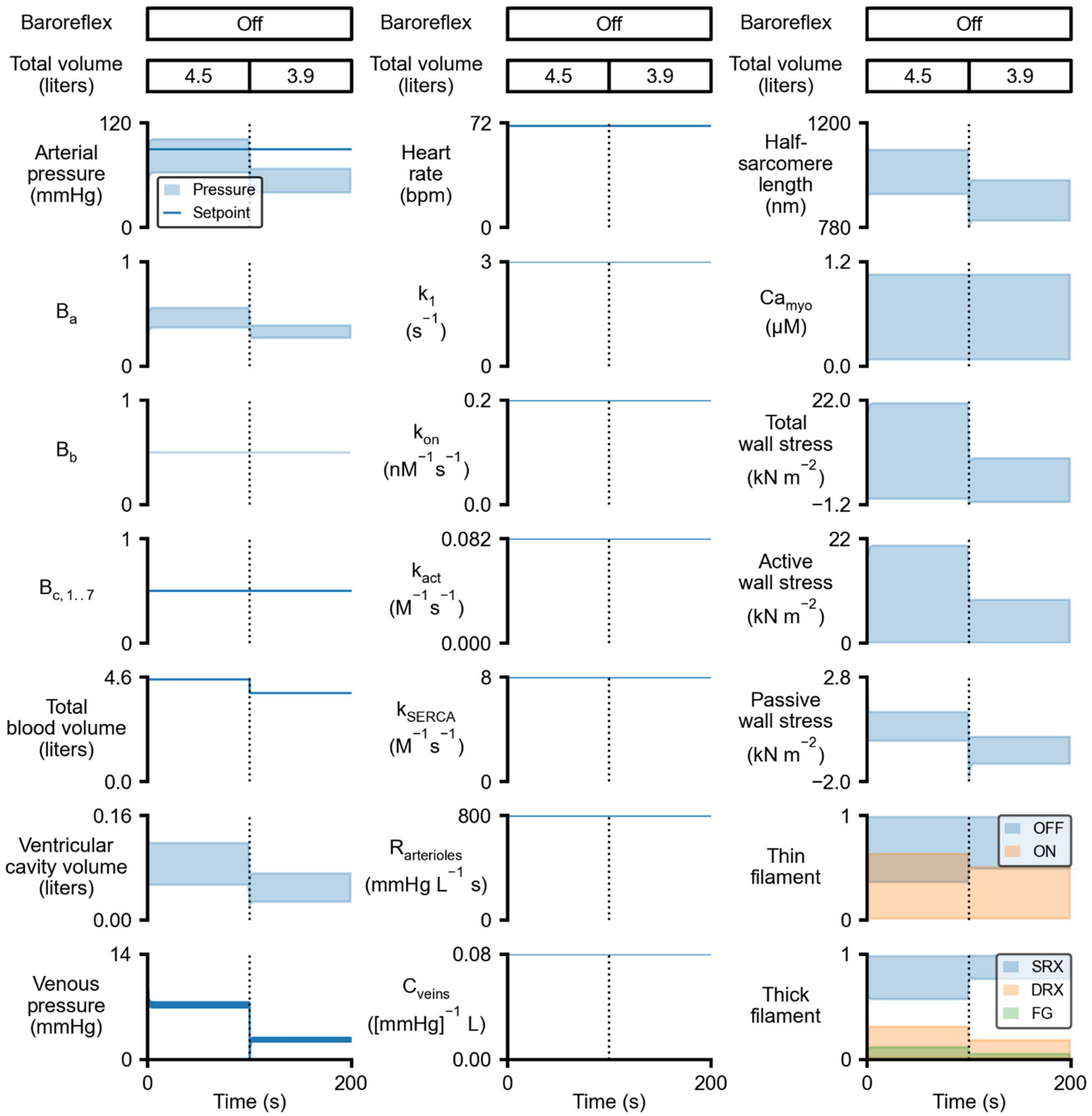


Fig S5: Simulated response to blood loss in the absence of baroreflex control.

This figure is identical to Fig 3 in the main text except that the baroreflex remains inactive. Arterial pressure falls to 67/40 mmHg when 600 ml of blood is removed from the venous compartment.

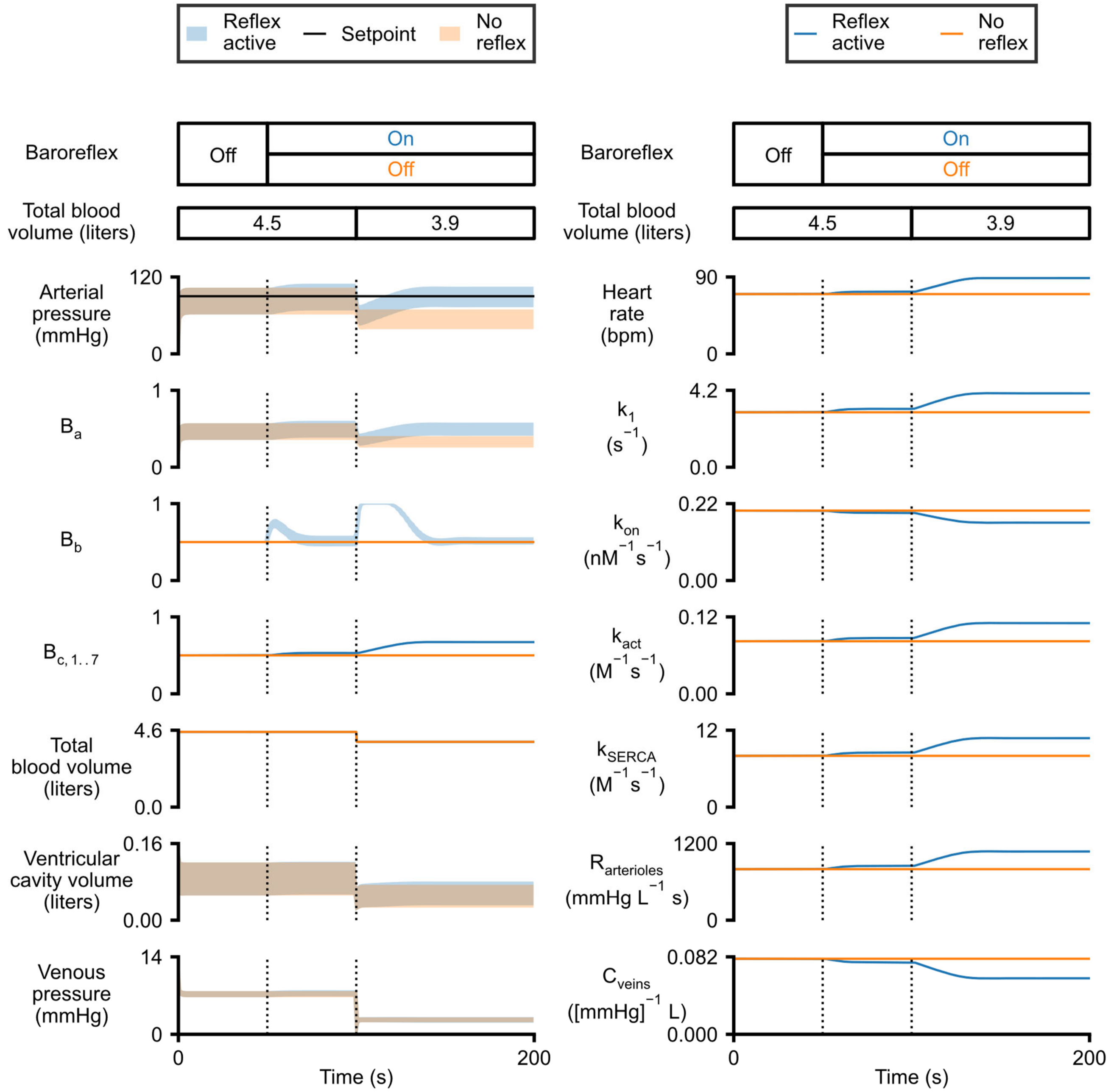


Fig S6: Simulations showing the response to blood loss with and without baroreflex control.

This figure replots data from Fig 3 in the main text and Fig S5 and demonstrates how the reflex algorithm returns arterial pressure to the original set-point after the total blood volume is reduced.

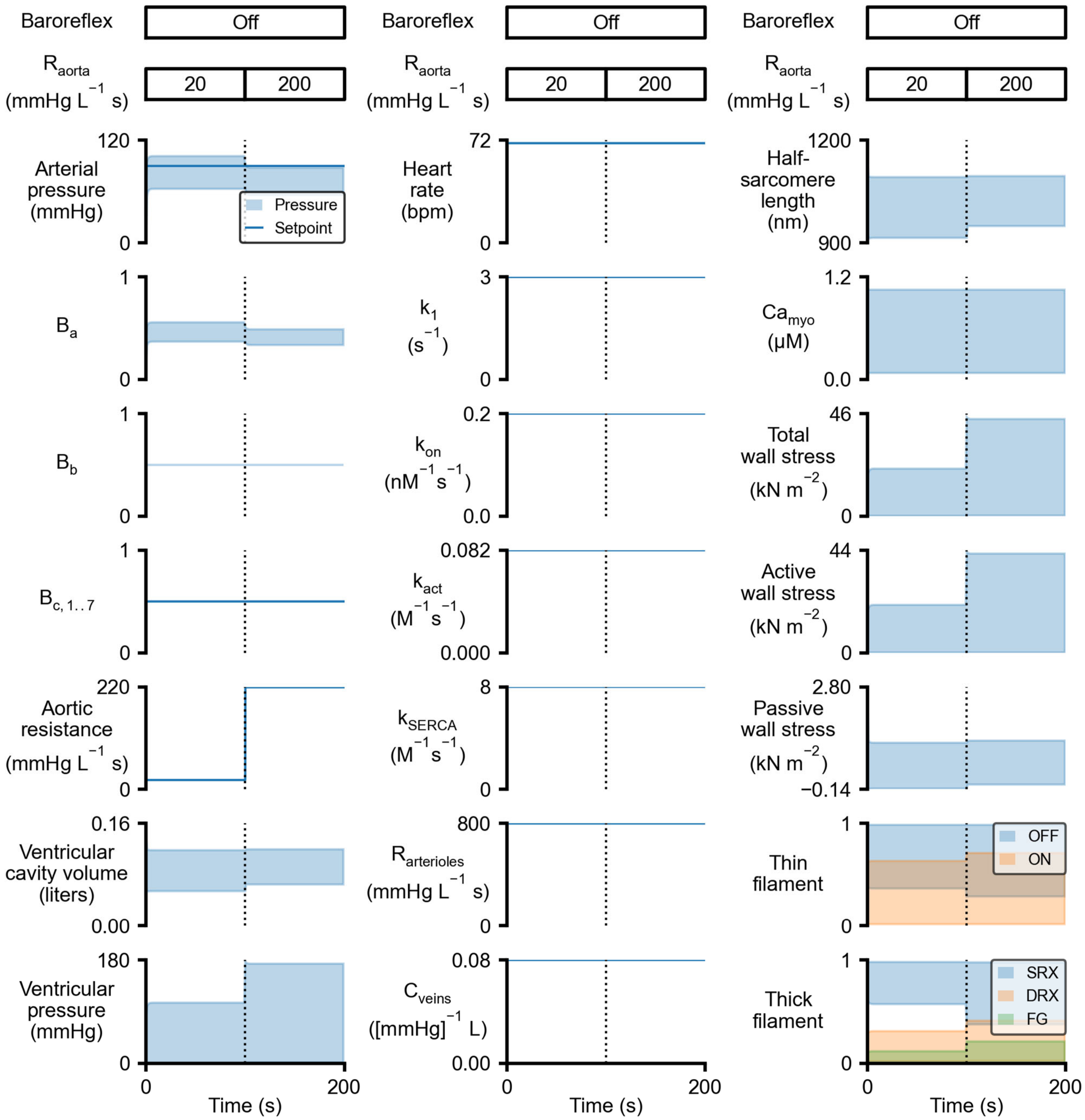


Fig S7: Simulated response to increased aortic resistance in the absence of baroreflex control.

This figure is identical to Fig 4 in the main text except that the baroreflex remains inactive. The mean arterial pressure falls from 84 to 73 mmHg when the aortic resistance is increased.

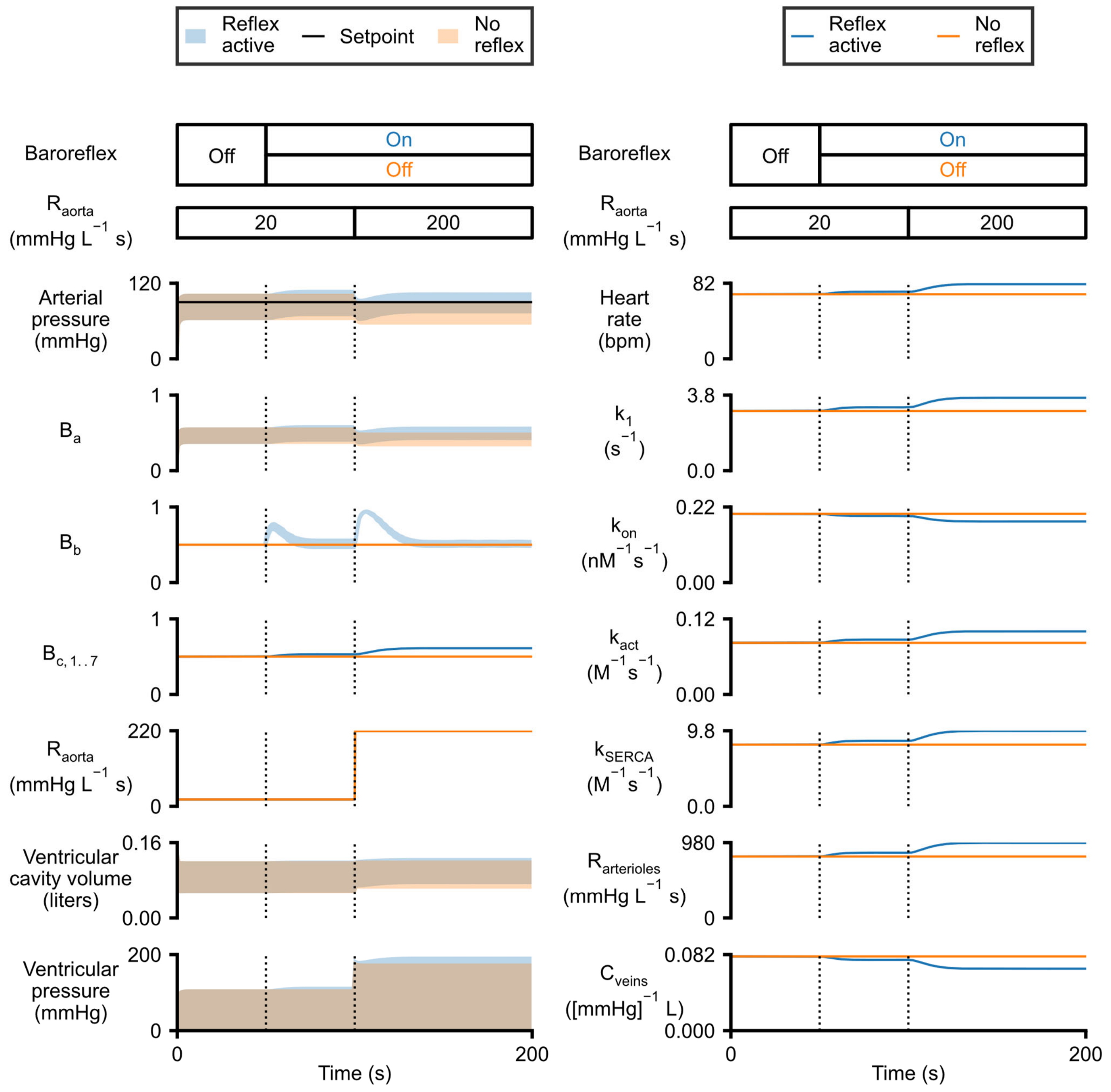


Fig S8: Simulations showing the response to increased aortic resistance with and without baroreflex control.

This figure replots data from Fig 4 in the main text and Fig S7 and demonstrates how the reflex algorithm returns arterial pressure to the original set-point after aortic resistance is increased.

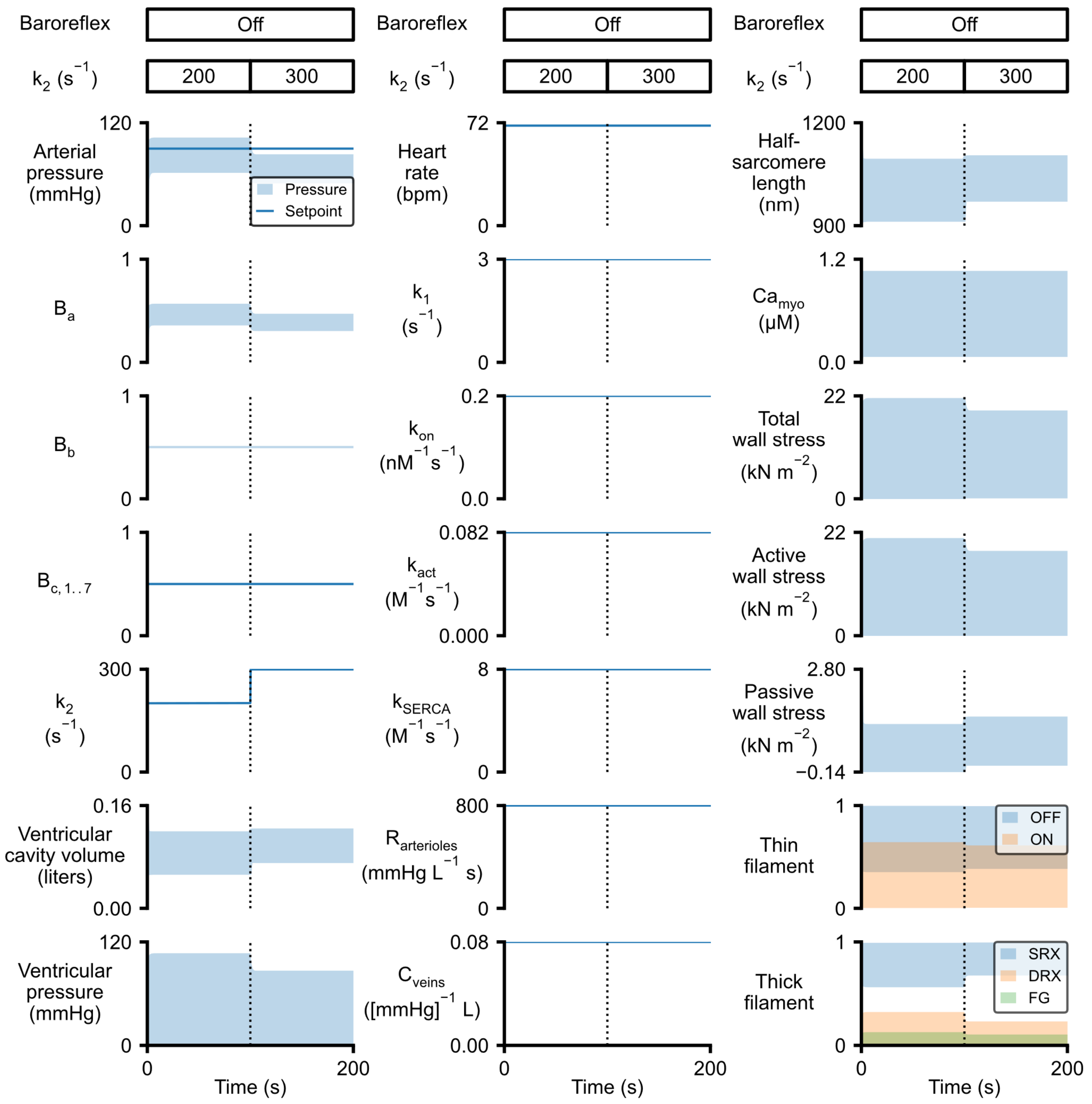


Fig S9: Simulated response to reduced myofilament contractility in the absence of baroreflex control.

This figure is identical to Fig 6 in the main text except that the baroreflex remains inactive. The mean arterial pressure falls from 83 to 67 mmHg when myosin heads are biased towards the super-relaxed SRX state.

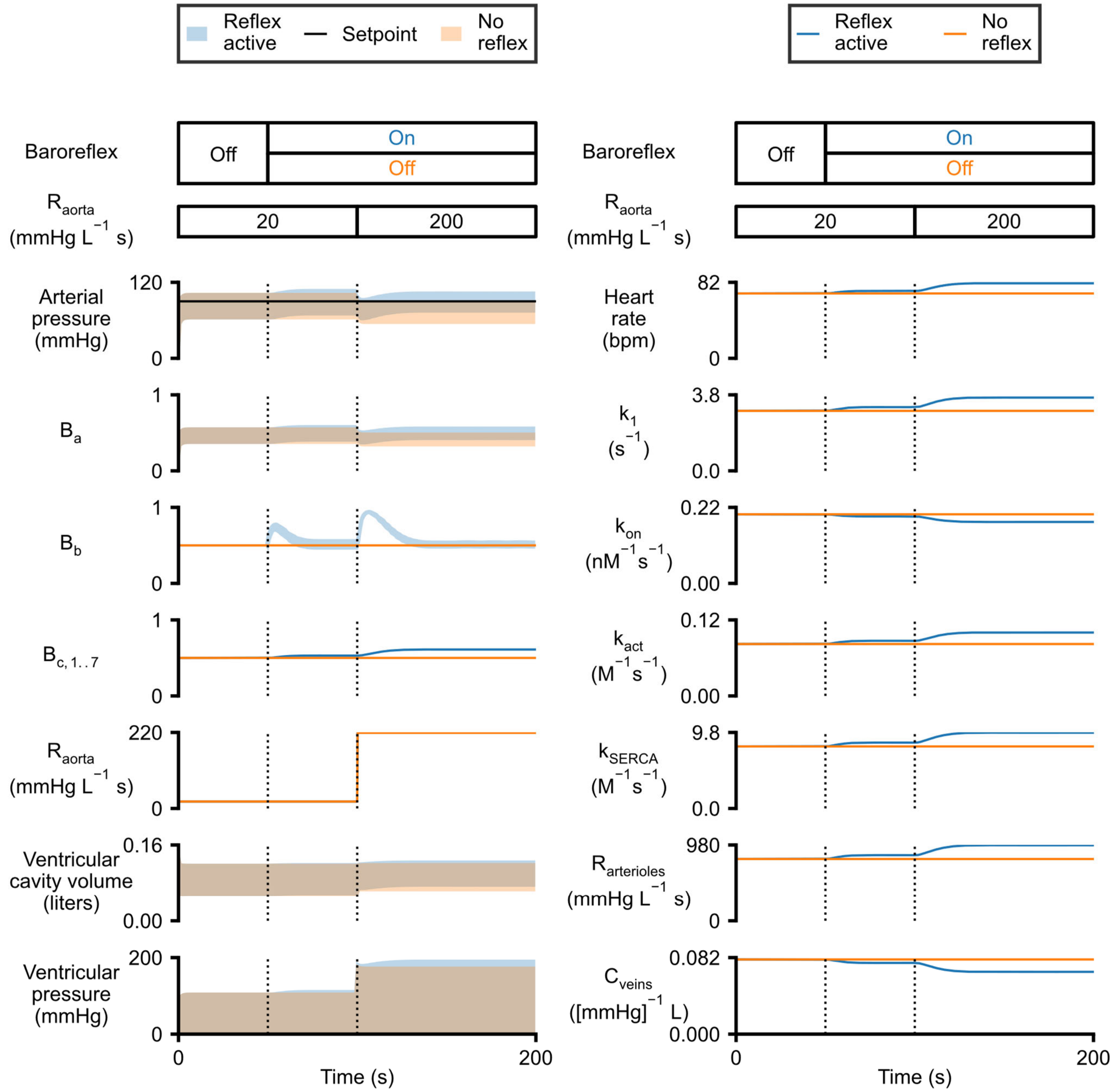


Fig S10: Simulations showing the response to reduced myofilament contractility with and without baroreflex control.

This figure replots data from Fig 6 in the main text and Fig S9 and demonstrates how the reflex algorithm returns arterial pressure to the original set-point after myosin heads are biased towards the super-relaxed SRX state.

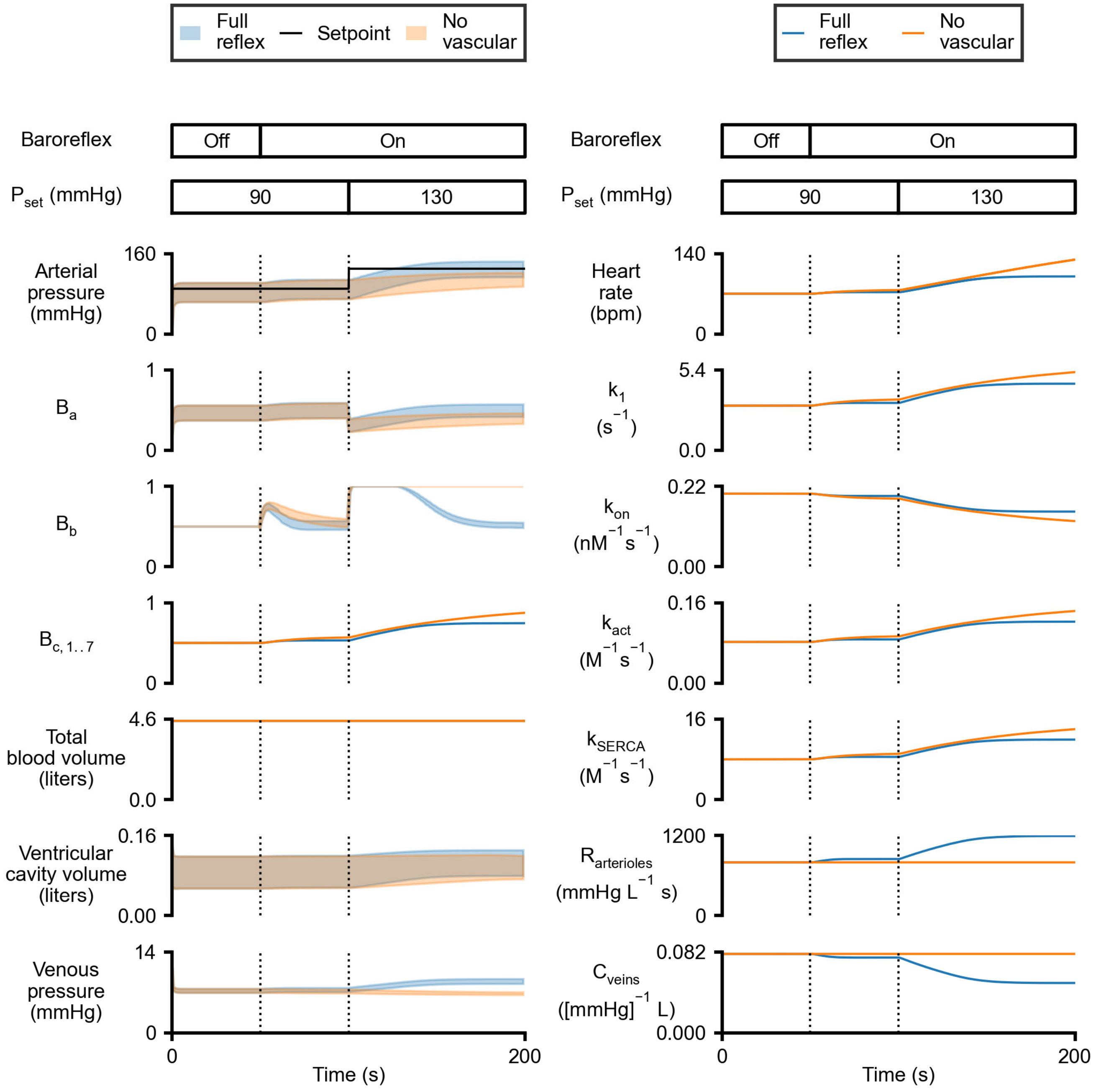


Fig S11: Simulations showing the response to a change in $P_{set}$ with and without baroreflex control of vascular tone.

This figure replots data from Figs 2 and 8 in the main text and demonstrates how including reflex control of arteriolar stiffness and venous compliance improves the speed and robustness of the reflex algorithm.
